# Isolation and characterization of a phage collection against *Pseudomonas putida*

**DOI:** 10.1101/2024.03.26.586772

**Authors:** Age Brauer, Sirli Rosendahl, Anu Kängsep, Alicja Cecylia Lewańczyk, Roger Rikberg, Rita Hõrak, Hedvig Tamman

**Affiliations:** Department of Bioinformatics, Institute of Molecular and Cell Biology, University of Tartu, Estonia; Department of Genetics, Institute of Molecular and Cell Biology, University of Tartu, Estonia

## Abstract

The environmental bacterium *Pseudomonas putida* has a wide range of metabolic pathways and thus entails great promise both as cell factories in biotechnological production and for bioremediation approaches to degrade various aromatic pollutants. To thrive in the environment, *P. putida* has to be able to endure the constant threat posed by bacteriophages. Surprisingly, up until now, only a few phages have been isolated for the common laboratory strain *P. putida* KT2440, and no phage defence mechanisms have been characterized yet. Here, we introduce a novel <u>C</u>ollection of <u>E</u>nvironmental *Pseudomonas putida* <u>P</u>hages from <u>Est</u>onia (CEPEST), consisting of 67 dsDNA phages that belong to 22 phage species and nine phage genera. We show that most of the phages in the CEPEST collection are more infectious at lower temperatures, have a narrow host range, and require intact LPS for *P. putida* infection. Additionally, we demonstrate that cryptic prophages in the *P. putida* chromosome strongly protect against the infection of many phages, whereas the chromosomal toxin-antitoxin systems play no role in phage defence of *P. putida*.

## Introduction

*Pseudomonas putida*, especially the strain KT2440 and its isogenic PaW85, has long served as a model species of environmental bacteria. Its main features include high stress tolerance and a wide variety of different metabolic pathways (Dos Santos *et al*., 2004; Reva *et al*., 2006; Nikel *et al*., 2016). The versatile metabolism of *P. putida* has increased the interest of the scientific and biotechnology communities in utilizing this microbe in bioproduction and biodegradation applications. To engineer a robust microbial cell factory from this laboratory work-horse, the genome of *P. putida* has been significantly edited and reduced (Martínez-García *et al*., 2014; Aparicio, De Lorenzo and Martínez-García, 2019; Martínez-García and De Lorenzo, 2024). Notably, while *P. putida* is considered to have high potential for bio-industry, it is curious that there is no knowledge regarding how this bacterium protects itself from its natural enemies, bacteriophages. Indeed, phages are abundant in the environment with the number of phage virions on Earth estimated to reach up to 10^31^ (Comeau *et al*., 2008). Lytic phages can efficiently kill their host by entering the cells, reprogramming the cell metabolism to produce viruses, and finally lysing the exhausted host bacterium to release the new progeny of viruses. Thus, phages pose a constant threat to both basic microbiology studies as well as bacteria-based bio-industrial applications (Bogosian, 2005; Los, 2012). Yet, despite several decades of research, the phage resistance of *P. putida* KT2440 or PaW85 has not been studied, and only a few phages able to infect the KT2440 strain have been isolated recently (Ngiam, Weynberg and Guo, 2022; Jaryenneh, Schoeniger and Mageeney, 2023).

Bacteria are endowed with different mechanisms that protect them against viruses. A vast number of phage defence systems operating through various ways have been revealed to the scientific community in recent years (Doron *et al*., 2018; Millman *et al*., 2022; Vassallo *et al*., 2022; Georjon and Bernheim, 2023). These systems are often encoded outside the core genomes of bacteria and tend to cluster in so-called defence islands of mobile DNA regions (Makarova *et al*., 2011). For example, many phage defence systems can be found in prophages, i.e., in genomes of temperate phages that have been integrated into the bacterial chromosome (Bondy-Denomy *et al*., 2016; Patel and Maxwell, 2023). Among the widespread anti-phage mechanisms are restriction-modification systems (Tock and Dryden, 2005), CRISPR-Cas modules (Horvath and Barrangou, 2010), and various types of abortive infection systems (Abi systems) (Lopatina, Tal and Sorek, 2020). Some Abi systems have been shown to act through a toxin-antitoxin mechanism (Bouchard *et al*., 2002; Fineran *et al*., 2009; Dy *et al*., 2014).

Toxin-antitoxin (TA) systems are widespread bacterial genetic elements that code for a potentially harmful toxin protein and an antidote, an RNA or a protein, able to inhibit the toxicity of its partner efficiently. TA systems are known to stabilize plasmids and other mobile elements (Ogura and Hiraga, 1983; Wozniak and Waldor, 2009; Fraikin and Van Melderen, 2024), but besides that, they have also been implicated in the phage defence of bacteria (LeRoux and Laub, 2022; Kelly *et al*., 2023). Phage infection has been shown to activate toxins. This, in turn, triggers the cell death or severe growth inhibition of phage-infected bacteria (Hazan and Engelberg-Kulka, 2004; Fineran *et al*., 2009; Dy *et al*., 2014; LeRoux *et al*., 2022), which limits the phage propagation within bacterial population. Upon activation, some toxins may even preferentially target phage products or processes (Guegler and Laub, 2021). Toxins can be activated either by phage-caused overall shutoff of host transcription, followed by destabilization of the RNA-protein TA complex (Short *et al*., 2018; Guegler and Laub, 2021) or by direct binding of a phage structural protein resulting in toxin liberation (Zhang *et al*., 2022). Still, despite accumulating evidence of TA systems acting as anti-phage elements, in most cases, the participation TA systems in phage defence has not been confirmed. Moreover, the activation mechanism of the toxin is often unidentified.

*P. putida* PaW85, as well as its isogenic KT2440 strain, harbours numerous chromosomal TA systems. Deletion of as many as 13 TA systems from the *P. putida* PaW85 genome (strain Δ13TA) revealed that multiple TA loci confer no clear fitness benefits but rather impose slight fitness costs to the bacterium, given that the presence of TA loci decreased the competitive fitness of wild-type *P. putida* (Rosendahl *et al*., 2020). Yet, so far, we have tested the effects of the lack of chromosomal TA loci on stress tolerance, persistence, biofilm formation, and competitive fitness (Rosendahl *et al*., 2020), but we have not yet analyzed whether the TA systems can protect *P. putida* from invading phages. As phage defence has been considered to be an important function of TA systems (Song and Wood, 2020; LeRoux and Laub, 2022; Kelly *et al*., 2023), it is reasonable to test the possibility that TA loci, despite their cost, are maintained in *P. putida* PaW85 chromosome due to their importance upon phage attacks.

Here, we present a collection of environmental bacteriophages isolated with a *P. putida* PaW85 derivative lacking 13 chromosomal TA loci and four cryptic prophages. Twenty-two phage species from nine genera were characterised by their genome sequences, taxonomy, morphology, temperature requirements, and host and receptor specificities. This first library of *P. putida* phages allowed us to test whether TA systems or prophages could be involved in the phage defence of *P. putida*. The collection opens up many research directions to follow, as the need to catch up on phage defence research of this bacterium is of crucial importance to the scientific and bio-industry communities.

## Experimental procedures

### Bacterial strains, media, and growth conditions

The bacterial strains and plasmids used are listed in Table S1 (Supplementary File 1). All strains constructed in this study are derivatives of *P. putida* PaW85 (Bayley *et al*., 1977), which is isogenic to well-studied KT2440 (Regenhardt *et al*., 2002). Bacteria were grown in lysogeny broth (LB). If selection was necessary, the growth medium was supplemented with kanamycin (50 µg ml^−1^) for *E. coli* and benzylpenicillin (1500 µg ml^−1^), kanamycin (50 µg ml^−1^) or streptomycin (200 µg ml^−1^) for *P. putida*. *E. coli* was incubated at 37 °C and *P. putida* at 30 °C, except for phage infection experiments that mainly were conducted at 20 °C. Bacteria were electrotransformed according to the protocol of Sharma and Schimke (Sharma and Schimke, 1996).

### Construction of strains

*P. putida* Δ4ϕ and Δ13TAΔ4ϕ strains were constructed by sequential deletion of four prophages from *P. putida* wild-type PaW85 and its Δ13TA derivative. The pEMG- and pSNW2-based plasmids were used for strain construction according to a well-described protocol (Martínez-García and de Lorenzo, 2011). Plasmids are listed in Table S1 (Supplementary File 1), and oligonucleotides used in PCR amplifications are listed in Table S2 (Supplementary File 1). Notably, first, the ΔP1 and the Δ13TAΔP1 strains devoid of prophage P1 were obtained by prophage spontaneous deletion when we aimed to delete the P1-encoded *hicAB2* (PP_3900-3899) toxin-antitoxin system by using plasmid pEMG-Δ*hicAB2*. However, instead of the *hicAB2* locus deletion, the whole P1 excised from the *P. putida* genome. The prophage P1 genome is surrounded by 67-bp-long direct repeats *attL* (genomic location 4371608-4371674) and *attR* (4427499-4427565) at each end, and sequencing of ΔP1 and the Δ13TAΔP1 strains revealed that recombination between *attL* and *attR* had created a clean P1 deletion leaving one *att* site. *P. putida* ΔP1 and Δ13TAΔP1 strains were used to delete other prophage genomes in the order P4, P3, and P2. The pEMG-based plasmids containing prophage deletion loci that were employed for strain generation were a generous gift from Esteban Martínez-García (Martínez-García *et al*., 2015). For the deletion of *wbpL* (PP_1804) gene, the upstream and downstream regions of *wbpL* were amplified separately with primer pairs del1804Eco/del1804 and del1804-pikk/del1804Bam, respectively. Two PCR products were joined into an approximately 1-kb fragment by overlap extension PCR using primer pair del1804Eco/del1804Bam, and inserted into EcoRI-BamHI opened plasmid pSNW2 (Volke *et al*., 2020), a *gfp-*containing derivative of pEMG.

### Isolation of bacteriophages

The enrichment method was used to isolate phages from various soil and water samples. The sampling date and exact source for each phage are indicated in Table S3 (Supplementary File 2). *P. putida* PaW85 Δ13TAΔ4ϕ strain was used as the host bacterium in the isolation process. To enrich the environmental samples with phages, 5 ml of 10x LB medium, 2 ml of exponential phase (OD_580_∼1) Δ13TAΔ4ϕ culture, CaCl_2_ (final concentration 10 mM), and ciprofloxacin (final concentration 0.01 µg/ml) were added to the 100 ml of environmental samples. Samples were incubated in flasks overnight at 70 rpm at 20 °C. The next day, to get rid of the bacteria and soil sediments, samples were centrifuged for 30 min at 2400 g, and the supernatant was filtrated (0.22 µm filter). To see if the phage isolation was successful, 1 ml of the filtrate was mixed with 200 µl of exponentially growing Δ13TAΔ4ϕ culture, and 5 ml of 0.3% melted LB agar (42 °C) containing 10 mM of CaCl_2_ and overlaid on LB plates supplemented with ciprofloxacin (0.03 µg/ml). Plates were incubated overnight at 20 °C. If plaques had formed, single plaques were isolated and purified. To do that, a single plaque was picked and suspended in 100 µl of SM buffer (50 mM Tris-HCl pH 7.5, 100 mM NaCl, 8 mM MgSO_4_, 0.01% gelatin). 5 µl of chloroform was added to lyse the bacteria. Ten-fold dilutions of the phage suspension were made, and 2 µl drops were spotted on bacterial lawn plates. Plates were incubated overnight at 20 °C. Next, a new single plaque was picked, and the purification step was repeated three times in total. Purified phages were stored in phage storage buffer (SM buffer) both at 4 °C and −80 °C.

### Phage DNA purification

For phage DNA extraction, 1 to 2 ml of phage solution with the titer ranging from 10^8^ to 10^10^ PFU/ml was treated with 2 units/ml of DNase and 100 μg/ml RNase (Thermo Scientific), followed by the phage DNA extraction with Phage DNA Isolation Kit (Norgen Biotek) according to the protocol supplied by the manufacturer.

On some cases, the DNA yield from the kit was extremely low, and other measures had to be taken. For the jumbo phages of cluster G3, GeneJET Genomic DNA Purification Kit columns (Thermo Scientific) proved more efficient. The protocol of the phage DNA purification (Norgen Biotek) kit was followed until DNA binding when columns from the genome extraction kit (Thermo Scientific) were used instead of the phage kit columns for binding and elution of the DNA. In some other cases when the DNA yield was low (e.g. phages from genus clusters G6, G7 and G9), the phages were first precipitated from phage lysate with 80 mM ZnCl_2_ for 5 minutes at 37 °C, pelleted by centrifugation at 10 000 g for 1 min and then resuspended in 400 μl of TES buffer (0.1 M Tris pH 8.2, 0.1 M EDTA, 0.3% SDS). Phage capsids were digested by adding 4 μl of proteinase K (Thermo Scientific) and incubating for 1 h at 60 °C, after which the proteins were precipitated with 40 μl 3 M K-acetate (pH 4.8) on ice for 15 minutes. The pelleted proteins were removed by centrifugation (1 min, 10 000 g at 4 °C) and DNA in the supernatant was pelleted by adding a 1:1 volume of isopropanol to the liquid fraction and incubated on ice for 5 minutes. The pellet of DNA was centrifuged down (10 000 g for 10 min), washed twice with 75% ethanol, and resuspended in 60-70 μl of water. To purify the precipitated DNA sample, it was cleaned with a DNA Clean-up and Concentration kit (Zymo Research) according to the manufacturer’s protocol and eluted with 35 μl of water. Regardless of the method used, all the purified phage DNA samples were sequenced.

### Sequencing and assembly

Whole genome sequencing was done with Illumina MiSeq PE 2 x 175 bp or PE 2 x 151 bp setup. The average sequencing depth was 120x. Sequencing reads were filtered with fastp v.0.21.0 (Chen *et al*., 2018) and assembled with SPAdes v.3.15.4 using isolate mode (Prjibelski *et al*., 2020). Genome completeness was confirmed by checking the existence of technical repeats in both contig ends with an exact length of the largest k-mer used during the assembly process by SPAdes and one repeat copy was removed.

### Genome annotation and characterization

tRNA and protein coding genes were annotated using Prokka v.1.14.6 (Seemann, 2014) together with PHROGs (Terzian *et al*., 2021). Overlaps between ORFs and tRNAs were allowed. Average GC% and coding potential were calculated for each genome. Assembled contigs were reordered for easier visualization of genomic comparisons. Phages with a close relative already available in the databases were reordered similarly to the published genome. For new species and genera, we used PhageTerm v.1.0.12 (Garneau *et al*., 2017) to predict packaging strategy and termini. If sequencing depth and library method allowed reliable PhageTerm prediction, the same predicted termini were used for other same species phages (identity >95%) as well. Genomes without reliable PhageTerm prediction in any of the close relatives were reordered by setting a large terminase subunit as the first gene if there was no overlap with the preceding ORF. In case of overlaps, arbitrary non-intragenic start was chosen but similar start was ensured in the isolates belonging to the same species or in the same genus if possible.

### Taxonomical clustering analyses

For the comparison to available published phage genomes, we used the monthly updated INPHARED database (Cook *et al*., 2021) (version date 1 Feb 2024) including ∼27 000 complete or near complete phage genomes available in GenBank. To limit the number of genomes for further similarity analysis, vCONTACT was used (Bin Jang *et al*., 2019) to select all possible close relatives from Inphared that cluster together with *P. putida* phages in sequenced library. All those close relative candidates were added to 68 library phages for VIRIDIC analysis to find genome wide similarities in order to determine the species and genera classification (genome sequence identity over 95% and 70% correspondingly). In case of borderline species identity (close to 95%), further detailed comparisons regarding the gene content and locations were made using Mauve (Darling *et al*., 2004) to align and visualize the annotated genomes. Whole genome coding sequence comparisons were visualized with clinker (Gilchrist and Chooi, 2021).

### Phylogenetic tree

Phylogenetic trees were based on the Muscle (Edgar, 2004) alignment of four concatenated protein (22 representative species) or gene sequences (all 67 phages): major head protein, DNA primase, spanin and terminase large subunit. Trees were calculated with iqtree v.1.6.12 (Nguyen *et al*., 2015) using ModelFinder (Kalyaanamoorthy *et al*., 2017) and 1000 bootstrap replicates and visualized in iTOL web application (Letunic and Bork, 2021).

### Transmission electron microscopy

For morphology analysis by transmission electron microscopy (TEM), the phage samples were concentrated by polyethylene glycol (PEG) precipitation. One ml of phage filtrate (∼10^9^ PFU/ml) was treated with 5.3% PEG 8000 and 0.33 M NaCl. After 1.5 h incubation at 4 °C, the phage particles were pelleted at 8000 g for 10 min at 4 °C. The phage precipitate was allowed to resuspend slowly in 20 μl SM buffer overnight at 4 °C. 10 μl of high-titer phage suspension (∼10^11^ PFU/ml) was incubated for 5 min on a formvar/carbon-coated grid. After the excess liquid was removed with filter paper, negative staining of phage samples was performed for 2-3 min with an aqueous solution of 2% phosphotungstic acid or 2% uranyl acetate. Excess liquid was removed, the grids were rinsed in demineralized water, and air-dried. The TEM images were taken using a Tecnai G2 Spirit BioTwin transmission electron microscope at a 120 kV accelerating voltage, and images were captured using an Orius SC1000 camera.

### Plaque assays

Bacteria were grown in LB medium overnight at 20 °C. The bacterial cultures were diluted 15-fold into fresh LB medium and grown until OD_580_∼1 at 20 °C. Next, 200 µl of the bacterial culture was mixed with 5 ml of melted 0.3% LB agar medium (42 °C) containing 10 mM CaCl_2_ and overlaid on 1.5% LB agar plates containing 0.03 µg/ml or 0.01 µg/ml ciprofloxacin to create a bacterial lawn. For qualitative phage resistance assay, 1.5 µl drops of phage lysates with the maximum titer were spotted on the bacterial lawn plates. Plates were incubated overnight at 20 °C. The formation of plaques was assessed.

To quantify the infectivity of phages, phage lysates were standardized to a titer of 10^7^–10^8^ PFU/ml. 1.5 µl drops of ten-fold dilutions of phage lysates were spotted on the bacterial lawns. Plates were incubated overnight at 20 °C. The number of plaques formed was counted to calculate the efficiency of plating in plaque-forming units per milliliter.

## Results

### Phage isolation and genome sequencing

To address the question of the importance of chromosomal TA systems in *P. putida* PaW85 phage defence, we started with isolating phages from different environmental samples. To rule out the possibility of TA systems protecting the cells from phage infection and thus inhibiting phage isolation, we decided to use the *P. putida* PaW85 strain lacking 13 genomic TA systems (Rosendahl *et al*., 2020). Also, to isolate a diverse phage collection of *Pseudomonas putida* phages, we needed as sensitive *P. putida* derivative as possible. While nothing was previously known about the phage resistance of *P. putida* PaW85, it is commonly well established that genomic prophages often carry phage-defence elements (Bondy-Denomy *et al*., 2016; Dedrick *et al*., 2017; Patel and Maxwell, 2023). *P. putida* PaW85 genome harbours four cryptic prophages (Martínez-García *et al*., 2015), and presuming that they may contribute to phage resistance, we deleted all these prophages from the strain that already lacked 13 TA systems. Thus, as a host for phage isolation, we used *P. putida* strain Δ13TAΔ4ϕ (Figure 1).

**Figure 1.**
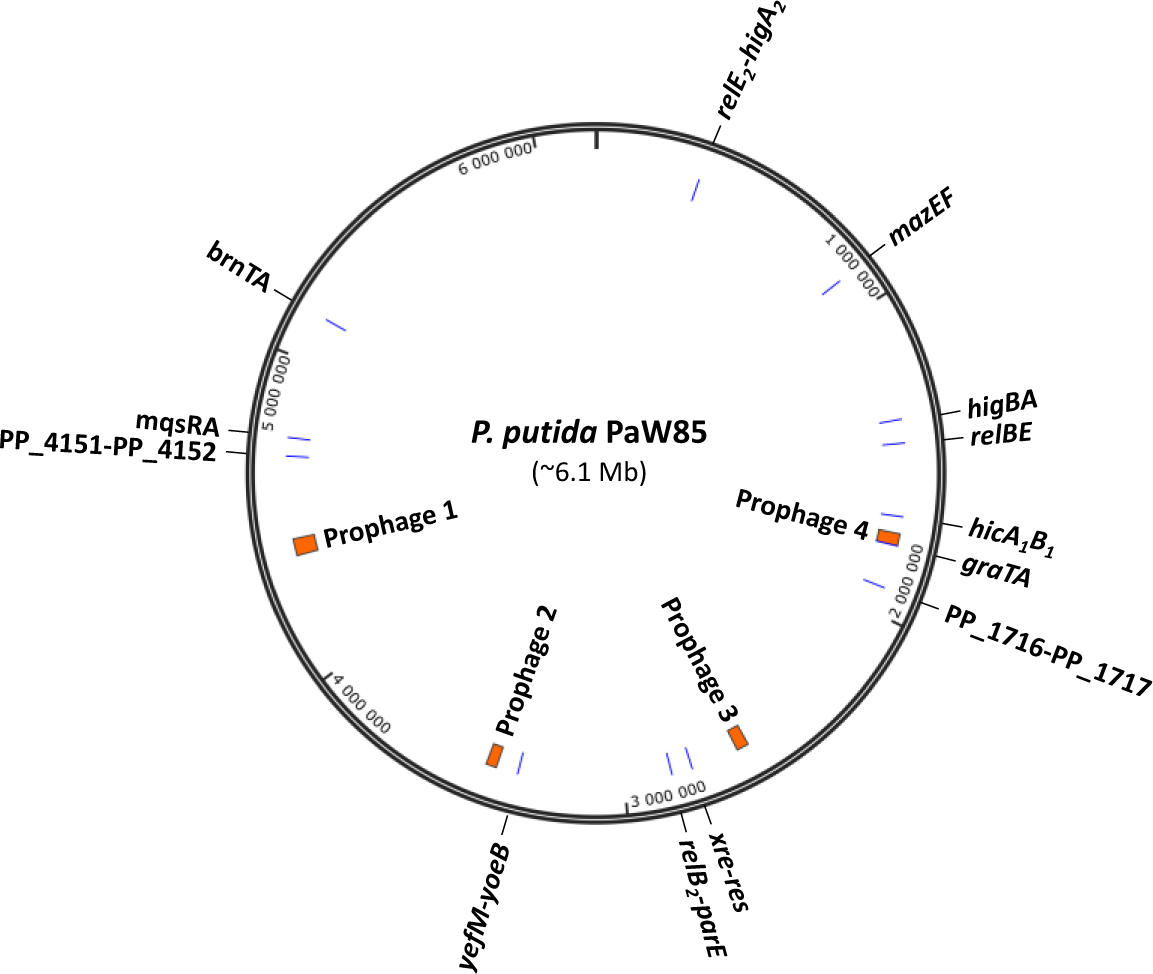
Schematic representation of *P. putida* PaW85 genome. The location of 13 toxin-antitoxin systems and four prophages that are deleted from the phage isolation strain *P. putida* Δ13TAΔ4ϕ are indicated. The figure was made with SnapGene software (www.snapgene.com).

Our first isolation attempts at the optimal growth temperature for *P. putida* of 30 °C were unsuccessful. Thus, we optimized the phage isolation protocol by lowering the incubation temperature. This was motivated by the fact that *P. putida* is an environmental bacterium that usually lives in the soil and water at much lower temperatures than 30 °C. As expected, reducing the temperature of the phage isolation procedure to 20 °C was fruitful and allowed us to collect 67 phages from different environmental samples (Table S3, Supplementary File 2). The phage collection was named CEPEST from the <u>C</u>ollection of <u>E</u>nvironmental Pseudomonas putida <u>P</u>hages from <u>Est</u>onia.

All isolated phages were sequenced, successfully assembled into complete genome sequences, and annotated. VIRIDIC (Moraru, Varsani and Kropinski, 2020) clustering analysis based on sequence similarity of assembled genomes demonstrated that 67 isolated phages belong to nine different genus clusters G1-G9 (Figure 2A; Figure S1, Supplementary File 3). Most sequenced genomes were 39-42 kb long, which is the most common length range among published phages (Figure S2, Supplementary File 3), and contained 45-52 open reading frames and no tRNA genes. Two genus clusters differed regarding the genome size. Cluster G1 phages with genome sizes 95-97 kb belonged to the size class that has so far been poorly represented among sequenced phages (Figure S2, Supplementary File 3). G1 genomes contained 169-176 protein-coding sequences and 16 to 17 tRNAs. Cluster G3 included eight phages with a genome length over 200 kb and were thus defined as jumbo phages.

**Figure 2.**
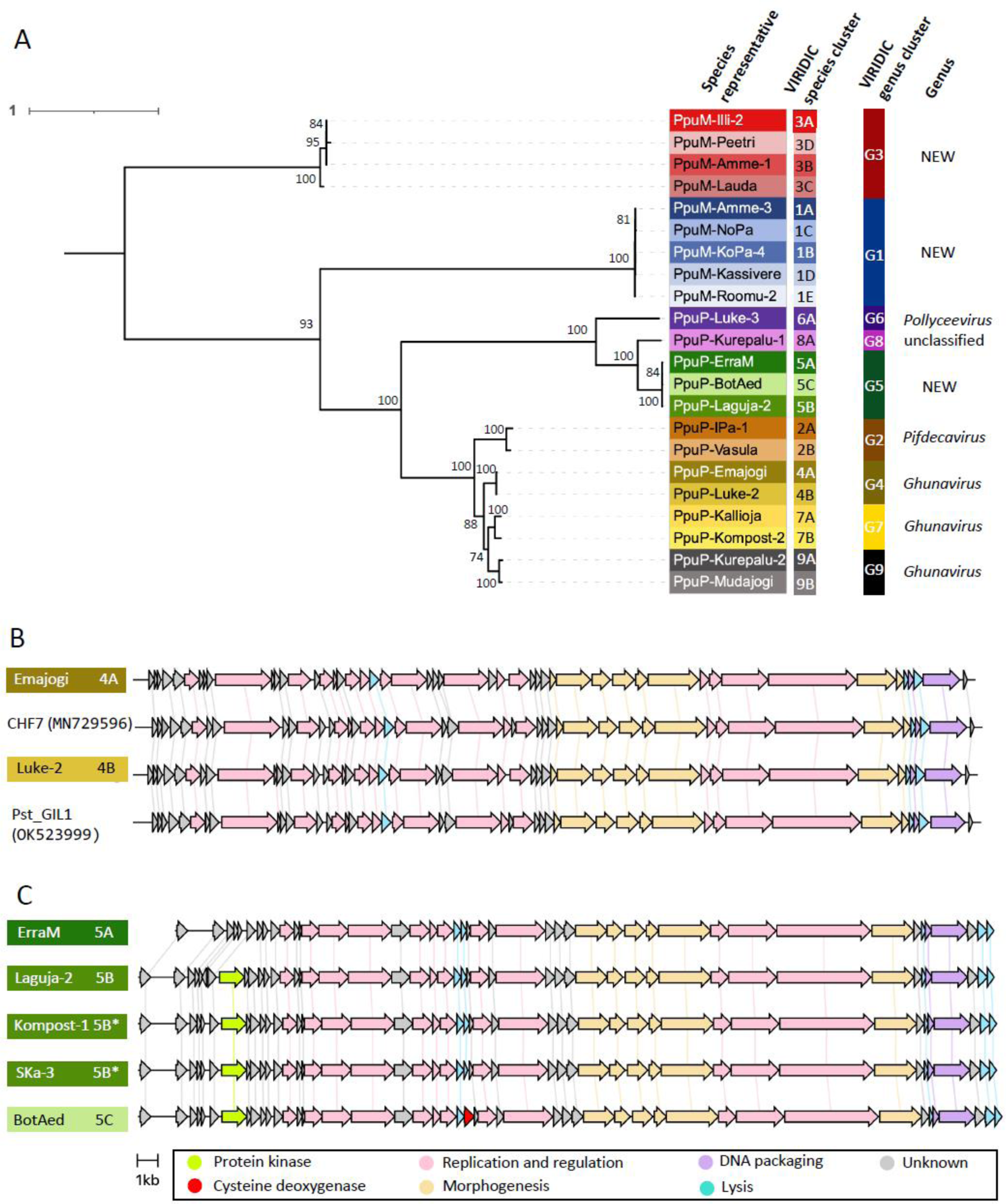
The phylogenetic analysis of phages in CEPEST collection. A: Midpoint-rooted phylogenetic tree of 22 representative phage species from nine genus clusters (G1-G9). The tree is based on multiple alignment of four concatenated protein sequences (major head protein, DNA primase, spanin, and terminase large subunit), calculated with IQ-TREE using ModelFinder and visualized using iTOL online tool. Bootstrap values >70 are shown. B: Gene content and organization in genus cluster G4 compared to published *P. putida* phages CHF7 and Pst_GIL1. C: Gene content and organization in genus cluster G5 species describing differences in gene content.

Genomic GC-content varied from 47.2 to 61.5%, being the lowest in G1 phages and highest in a G8 phage. All informative statistics for each sequenced genome are presented in Table S3 (Supplementary File 2).

### Phage phylogeny, taxonomy, and diversity

We compared all 67 sequenced genomes to complete phage genomes in GenBank provided in Inphared collection (Cook *et al*., 2021) as of February 1^st^ 2024, to determine taxonomical affiliation and distribution.

We were able to classify phages in cluster G2 as members of the genus *Pifdecavirus;* phages in G4, G7, and G9 as members of the genus *Ghunaviru*s; and phages in G6 belonged to the genus *Pollyceevirus* (Tables S7, S10, and S12, Supplementary File 2). A phage belonging to cluster G8 had a low genus level similarity (∼72%) to two *P. putida* phages LNA8 and LNA10 which are currently taxonomically unclassified (Ngiam, Weynberg and Guo, 2022) (Table S11, Supplementary File 2). Phages in three genus clusters (G1, G3, and G5) did not reach genus-level similarity (70%) to any of the available phage genomes, suggesting three new genera of phages (Tables S4-S6 and S8, Supplementary File 2).

Only two phages belonging to cluster G4 (*Ghunavirus*) had species level relatives among the Inphared collection. Phage Emajogi showed >95% similarity to Pseudomonas phages CHF7 (MN729596), phiPSA2 (KJ507099), and phiPsa17 (KR091952). Phage Luke-2 had >96% similarity to Pseudomonas phage Pst_GIL1 (OK523999) and >95% similarity to Pseudomonas phage KNP (KY798121) (Figure 2B; Table S7, Supplementary File 2).

Jumbo phage cluster G3 contained one phage named Aura with a genome size just below the jumbo phage length threshold (199 504 bp) but showed high similarity on genome sequence level and was therefore assigned to species cluster 3A together with Illi-2 and SKa-4 jumbo phages (Table S6, Supplementary File 2).

Genus clusters G2 and G5 were the most abundantly represented in our collection (Figure S1, Supplementary File 3). Majority of the isolates in both of those clusters were assigned to one species based on a comparison of genome sequences, but the third new genus cluster G5 needed additional analysis to decide the species assignment. There seemed to be three separate species in G5, but the final division remained ambiguous for two phages – SKa-3 and Kompost-1. VIRIDIC automatic clustering grouped these phages together with phage BotAed (species cluster 5C). However, the similarity matrix showed SKa-3 and Kompost-1 being more closely related to phages in species cluster 5B (Table S8, Supplementary File 2). In addition, BotAed contains a gene for cysteine dioxygenase, which is not present in other G5 phages, including SKa-3 and Kompost-1 (Figure 2C). Therefore, SKa-3 and Kompost-1 were marked as species 5B* (Table S8, Supplementary File 2).

Gene content analysis was also made for phage Vanda in G5. Although Vanda had genomic similarity reaching over 95% species level threshold with some 5B phages (Table S8, Supplementary File 2), we still marked Vanda as a 5A species phage. Namely, Vanda lacked a protein kinase coding gene similar to 5A phages ErraM (Figure 2C) and ErraS in contrast to 5B or 5C phages. Vanda belonging to 5A was also supported by VIRIDIC automatic species clustering.

In conclusion, VIRIDIC clustering, sequence similarity matrix, gene content, and organization comparison resulted in 22 representative species among nine identified genus clusters (Figure 2A). Twenty of those were new species without close and characterized relatives among publicly available bacteriophage genomes. Therefore, to know more about those phages, we set out to determine the morphology of the isolated phages with transmission electron microscopy (TEM). At least one representative species from each of the nine phage genera was analysed. TEM results demonstrated that most of the phages in our collection had podovirus morphology, and only phages from genus clusters G1 and G3 represented myoviruses (Figure 3; Table S3, Supplementary File 2). All the phages with podovirus virions share a similar genome and capsid size, around 40 kbp and 50-60 nm in diameter, respectively. Myoviruses from genus clusters G1 and G3 that have significantly larger genomes (almost 100 kbp for G1 and over 200 kbp for G3) than podoviruses also have larger capsids (Figure 3). For instance, the jumbo phages from genus cluster G3 have the largest virions of the collection with a head diameter of almost 100 nm. Thus, the morphological studies demonstrate that the genome sizes correlate with the morphology of the phages, with the two myovirus genera having larger genomes and larger virions than the podovirus phages in our collection.

**Figure 3.**
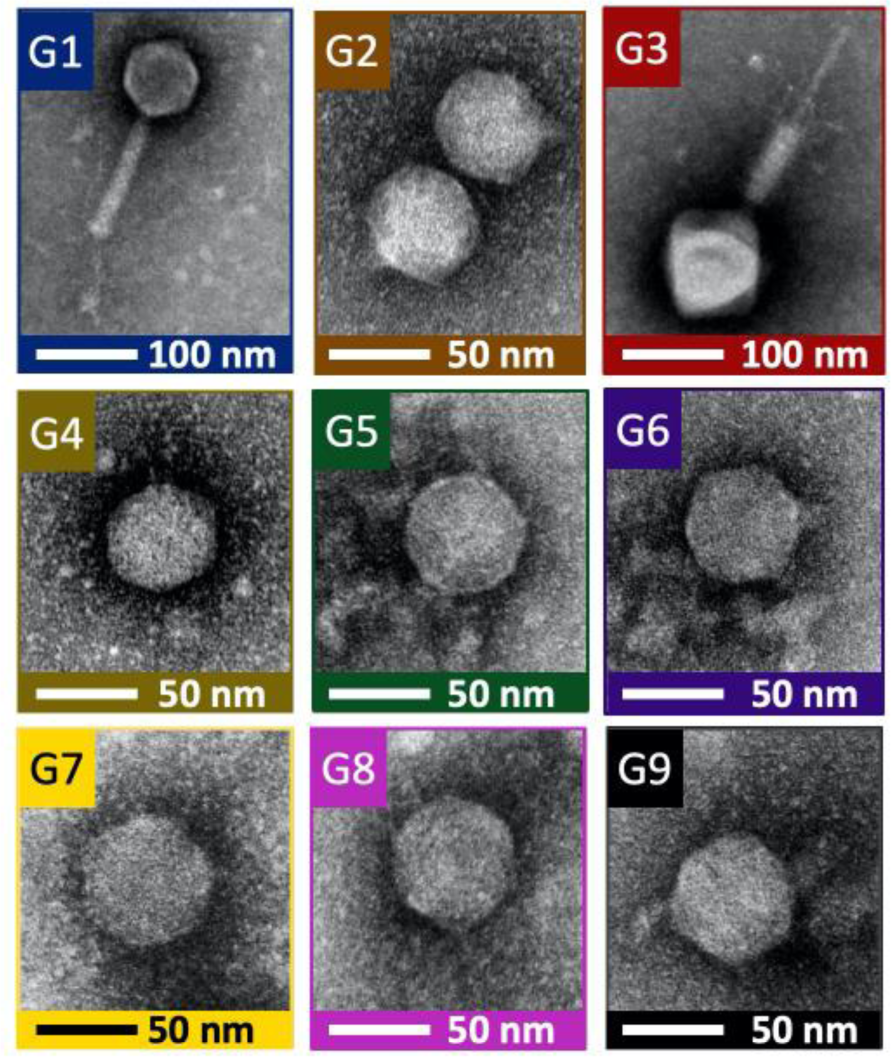
Morphology of phages in CEPEST collection. A negative staining TEM image of a representative phage from each genus cluster in the CEPEST collection (G1 – NoPa, G2 – Vasula, G3 – Amme-1, G4 – Emajogi, G5 – Laguja-2, G6 – Luke-3, G7 – Kompost-2, G8 – Kurepalu-1 and G9 – Kurepalu-2).

### Temperature-sensitivity of *P. putida* phages

Given that our first attempts for phage sampling at 30 °C were unsuccessful and that the phage library was collected at 20 °C, we hypothesised that the infection of *P. putida* phages might depend on the temperature. To determine the temperature range and sensitivity of the phages, we measured their infection ability at various temperatures, ranging from 15 to 37 °C. The results showed that most of the isolated phages are unable to infect the phage isolation host strain *P. putida* Δ13TAΔ4ϕ at 30 °C and 37 °C (Figure 4). Moreover, the infection efficiency of most phages was already strongly reduced at 25 °C, with the phages from genera G6 and G7 and the species 9B losing their infection ability already at 22 °C. However, all phages retained the ability to infect the *P. putida* cells at 15 °C, and, except for most genera G1 and G8 species, even presented the strongest infection phenotype at this low temperature (Figure 4). The most insensitive to changes in infection temperature were the phages from the genus cluster G1, as they (except for 1C phage NoPa) were infectious at all tested temperatures. Remarkably, the temperature sensitivity of phages can vary between species in one genus, e. g. phage 1C and most strongly 9B losing the ability to infect cells at clearly lower temperatures than other members of the corresponding genera (Figure 4; Figure S3, Supplementary File 3). Taken together, most *P. putida* phages in our collection are temperature-sensitive and lose their infection efficiency at higher than 20-25 °C.

**Figure 4:**
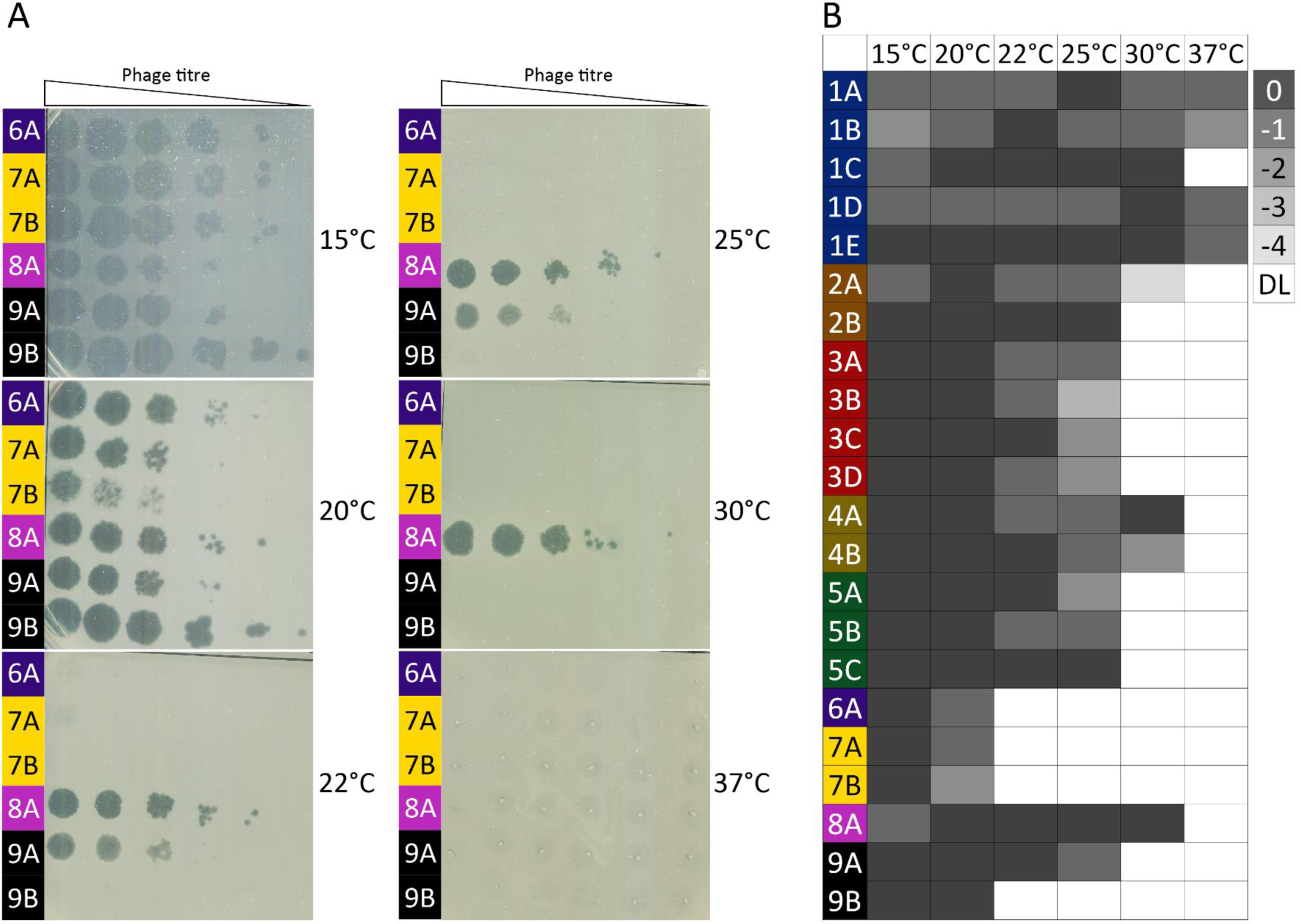
Temperature sensitivity of CEPEST phages. The infection efficiency of each phage species representative was tested at temperatures from 15 to 37 °C. A: Qualitative infectivity of phages from genus clusters G6, G7, G8, and G9. B: Heatmap of infection of EOP of each phage at a certain temperature. 0 level shows the most efficient infection, and numbers (and decreasing intensity of grey colour) represent a 10-time decrease in EOP. DL (white) is the detection limit of the experiment, no infection could be detected.

### Prophages, but not chromosomal toxin-antitoxin systems, increase the resistance of *P. putida* against several phages

To address our starting goal of studying the importance of 13 chromosomal TA systems in the phage defence of *P. putida* PaW85, we compared the phage resistance of the *P. putida* wild-type strain PaW85 and its Δ13TA derivative. Analysis of the infection of the 22 phages representing each species showed that the absence of TA systems did not affect the infection efficiency of phages (Figure 5A and B). Given that several phage species in our collection contain multiple phage isolates with slightly varying genome identities (Tables S3-S12, Supplementary File 2), we also compared the phage resistance of wild-type and Δ13TA with other 45 phages in our collection. Similar to the data obtained with 22 representatives of each species (Figure 5A and B), we recorded no difference between the phage sensitivity of the *P. putida* wild-type and Δ13TA strains (data not shown). Thus, the 13 chromosomal TA systems of *P. putida* do not function as phage defence modules against any of the phages in our collection.

**Figure 5:**
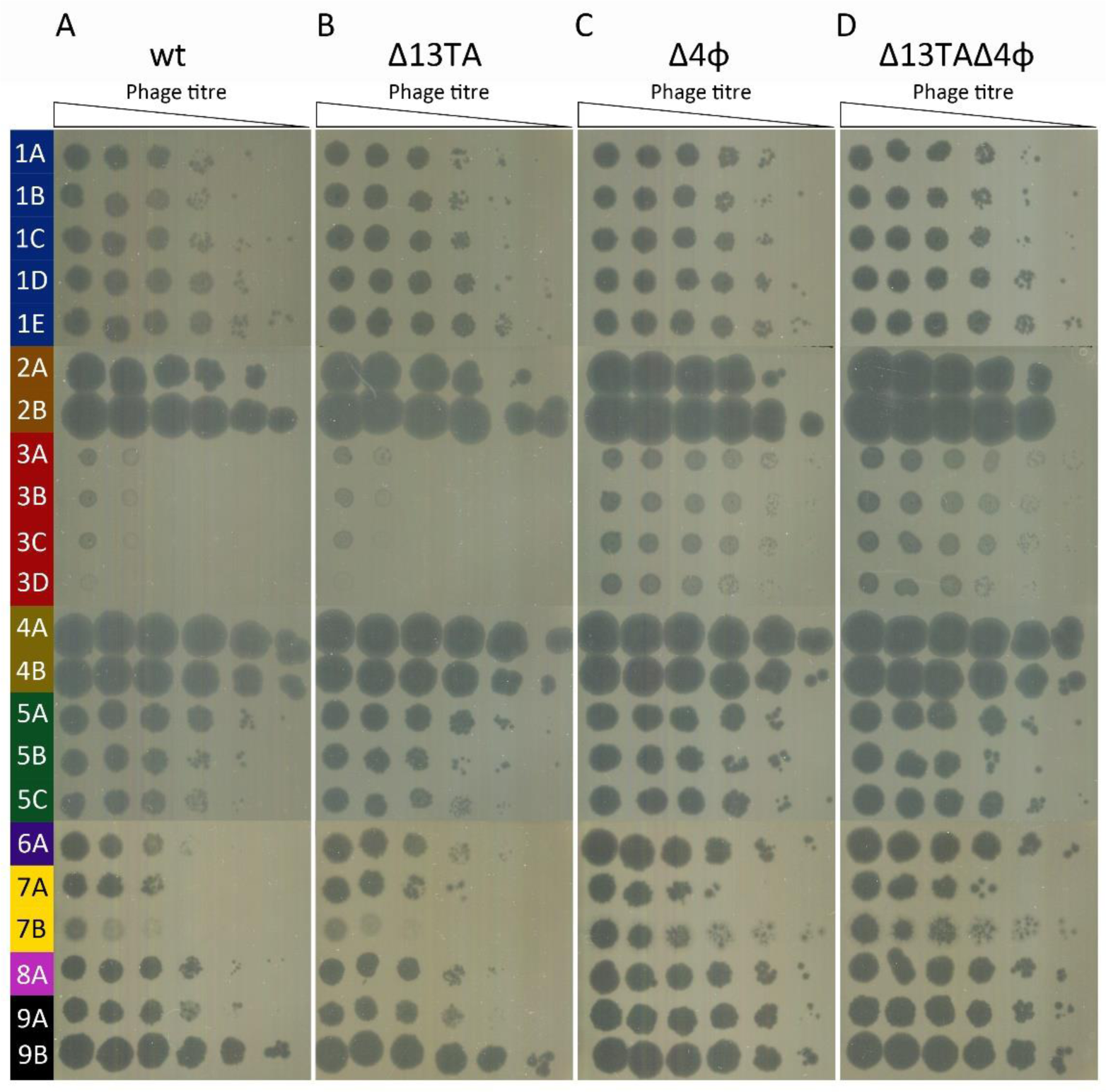
*P. putida* PaW85 prophages provide protection against several CEPEST collection phages. Quantitative EOP measurement of phage infection on bacterial lawns of A: *P. putida* PaW85 (wt), B: *P. putida* PaW85 lacking 13 TA systems (Δ13TA), C: *P. putida* PaW85 lacking four prophages (Δ4ϕ), D: *P. putida* PaW85 lacking 13 TA systems and four prophages (Δ13TAΔ4ϕ). 1.5 uL drops of 10-fold serial dilutions of phages were spotted on the strain to be tested, and EOP was counted from the plaques in dilution spots.

Interestingly, analysis of the wild-type and the Δ13TA strains revealed that the infection efficiency of some phages, particularly jumbo phages from genus G3, was very low against both strains (Figure 5A and B). This indicated that prophages that were present in wild-type and Δ13TA but were deleted from the phage isolation strain Δ13TAΔ4ϕ contribute to the phage resistance of *P. putida*. To test this, we compared the wild-type and the prophage-deficient Δ4ϕ strains and indeed observed that the latter was more sensitive to many phages (Figure 5A and C). The most prominent protective effect of prophages was observed against the G3 jumbo and 7B phages, as the wild-type strain demonstrated more than 1000-fold higher resistance than the Δ4ϕ strain (Figure 5A and C). Prophages also provided clear protection against the phage species 5C, 6A, 8A (about 10-fold), and 9A (about 100-fold). At the same time, infection of phages from genera G1, G2, G4, and the species 5A, 5B, and 9B was very little or not at all affected by the presence of prophages (Figure 5A and C). Also, the comparison of the phage resistance of Δ4ϕ and Δ13TAΔ4ϕ demonstrated similar phage sensitivity of the two strains (Figure 5C and D), which further confirms that 13 TA systems do not contribute to the phage resistance of *P. putida*. Overall, these results show that the four cryptic prophages in PaW85 provide a variable level of defence against about half of the phage species in the CEPEST collection. The rest of the phage species, however, are essentially insensitive to the presence of the prophages.

### *P. putida* phages in the CEPEST collection have a narrow host range

To study the host range of the isolated *P. putida* phages, we tested whether they can infect other bacteria from the genus *Pseudomonas*. To this end, we selected 11 different *P. putida* strains, eight strains of *P. syringae*, and two each of *P. fluorescens, P. aeruginosa,* and *P. stutzeri* (Table S1, Supplementary File 1) from the CELMS microbe collection of our institute (http://eemb.ut.ee/celms/main_list.php). As the closest relative to the *P. putida* wild-type strain PaW85, we also included its ancestor strain PaW1 (mt-2) that harbours the TOL plasmid pWW0 (Worsey and Williams, 1975). For the phage susceptibility testing of these 27 *Pseudomonas* strains, we used the highest phage concentrations of all 22 species in our collection (Figure 6A). The results show that the phages in the CEPEST collection are highly host-specific, with phages from genera G1, G5, and G8 being unable to infect any other strains apart from *P. putida* PaW85 and its ancestor mt-2 (Figure 6A and D). Only eight phage species out of 22 could infect some *P. putida* (strains CRTN6, PC13, and T9) and *P. syringae* (P82, DC3000, and B782a) strains. No phage was able to infect any of the tested *P. fluorescens, P. aeruginosa* or *P. stutzeri* strains (Figure 6A and D).

**Figure 6.**
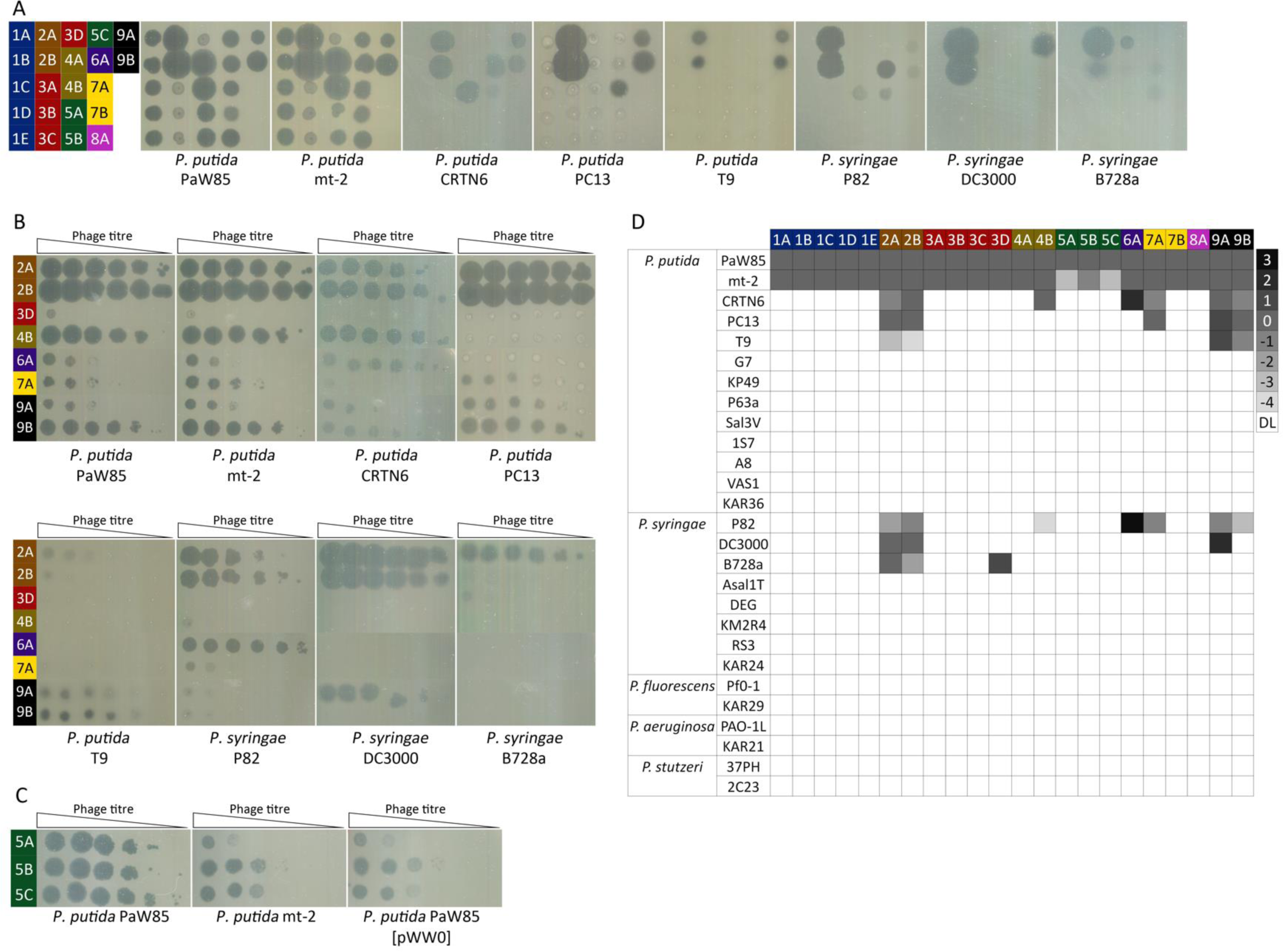
Host specificity of CEPEST phages. A: Qualitative assay of infection of all 22 phage species representatives (indicated in the colour matrix) on different *Pseudomonas* hosts. B: Quantitative EOP measurement of phage infection on their *Pseudomonas* hosts. 1.5 uL drops of 10-fold serial dilutions of phages were spotted on the strain to be tested, and EOP was counted from the plaques. C: EOP of G5 phages on *P. putida* PaW85, its ancestral strain mt-2 containing the pWW0 plasmid, and *P. putida* PaW85 with the pWW0 plasmid. D: A heatmap of host specificity of phages. 0 level shows the infection of PaW85 reference strain, and numbers (and decreasing or increasing intensity of grey colour) represent 10-fold changes in EOP. DL (white) is the detection limit of the experiment, no infection could be detected.

Out of the eight species of phages that can infect more strains than just the PaW background, the phages of G2 seem to have the widest host range, followed by 9A species. These phages infect not only five of the tested *P. putida* strains but also three (phages from genus G2) or two (phage 9A) *P. syringae* strains. The host range of phage species 3D, 4B, 6A, 7A, and 9B is narrower, but it is interesting to note that the same particular *P. putida* or *P. syringae* strains that were infected by G2 and 9A phages were also susceptible to some of these phages (Figure 6A and D).

To quantify the infection efficiency of 8 phages that could infect different *Pseudomonas* hosts, the phage resistance assay was carried out using serial dilutions of phages (Figure 6B). Results show that phages differ strongly in their ability to infect the tested hosts. Some phages, including 2A, 2B, 4B, 7A, or 9B, lyse some strains equally to *P. putida* PaW85, while they are less efficient against other susceptible hosts (Figure 6B and D). Surprisingly, 3D, 6A, and 9A phages could infect some *P. putida* or *P. syringae* strains even better than the *P. putida* PaW85 reference strain.

The phage sensitivity of *P. putida* PaW85 and its ancestor strain mt-2 seemed highly similar when the strains were infected with high phage titres (Figure 6A). However, infection efficiency quantification experiments revealed that the mt-2 strain was actually significantly more resistant to genus G5 phages than *P. putida* PaW85 (Figure 6C). The main difference between the two strains is that mt-2 contains the 117 kbp TOL plasmid pWW0. As plasmids have been shown to affect phage infection efficiency (Ngiam, Weynberg and Guo, 2022), we hypothesised that the TOL plasmid could be behind the protection against G5 phages. We tested a *P. putida* PaW85 that harbours the pWW0 plasmid (lab collection) and, indeed, all of the G5 phages infected this strain 10^2^ to 10^4^ times less efficiently (Figure 6C), showing that the plasmid pWW0 was causing the reduced infection efficiency of these phages. However, no other phage was affected by the presence of the TOL plasmid.

Interestingly, our results clearly show how two phage species from one genus can have completely different host specificity. Besides having variable infection efficiency towards the same strains, even the strain specificity may differ. For instance, the 4B and 7A species showed wider specificity: 4B could additionally infect *P. putida* strain CRTN6 and *P. syringae* P82, and 7A could infect *P. putida* PC13. At the same time, the other species in those genera, 4A and 7B, could not infect any cells except the PaW background (Figure 6A). Even more remarkably, phage 9A lysed the *P. syringae* strain DC3000 100 times more efficiently than the reference strain *P. putida* PaW85, while phage 9B could not infect that *P. syringae* strain at all (Figure 6A, B, and D). Taken together, 14 out of the 22 species of phages tested can only infect the *P. putida* PaW85 (and its predecessor and derivative strains), showing a very narrow host range for these phages.

### Most CEPEST collection phages need intact LPS for *P. putida* infection

As many podo- and myoviruses of Gram-negative bacteria have been shown to use polysaccharide sugar moieties for adsorption (Bertozzi Silva, Storms and Sauvageau, 2016), we set out to test whether the LPS of *P. putida* could be the receptor for phage adsorption of *P. putida* phages from our collection. As many phages recognize the outermost O-antigen part of LPS (Lindberg, 1973; Nobrega *et al*., 2018), we deleted the *wbpL* glycosyltransferase gene that is required for O-antigen synthesis (Rocchetta *et al*., 1998) from *P. putida* PaW85 wild-type and also from its Δ13TAΔ4ϕ derivative. The latter strain was constructed to test the receptor dependence of the G3 jumbo phages that infect the wild-type bacteria very poorly. We also picked a *P. putida* transposon mutant from our lab strain collection with a disrupted *wbpM* gene. Similarly to WbpL, WbpM is also involved at the beginning of the LPS O-antigen synthesis pathway (Creuzenet and Lam, 2001), and its disruption results in LPS molecules deprived of O-antigen moieties (Bélanger, Burrows and Lam, 1999). Analysis of these strains showed that out of the 22 phages, 18 could not infect the strains with either the *wbpL* deletion or *wbpM* truncation (Figure 7A). The only phages that retained the ability to infect LPS-deficient *P. putida* derivatives were two species from genus G2 and phages 4A and 7B. These four phages were further analysed to quantify their infection efficiency against O-antigen-deficient strains. Surprisingly, the deficient LPS synthesis significantly increased the infection efficiency of 7B phage Kompost-2, where the ΔwbpL strain was about 1000-fold more sensitive than the wild-type strain (Figure 7B). For the other three phages (2A, 2B, and 4A), the infection efficiency against the LPS mutants remained unchanged (Figure 7B). These results confirm that the LPS, particularly its outermost O-antigen moiety, is the main receptor for most of the 22 species of phages in our CEPEST collection.

**Figure 7:**
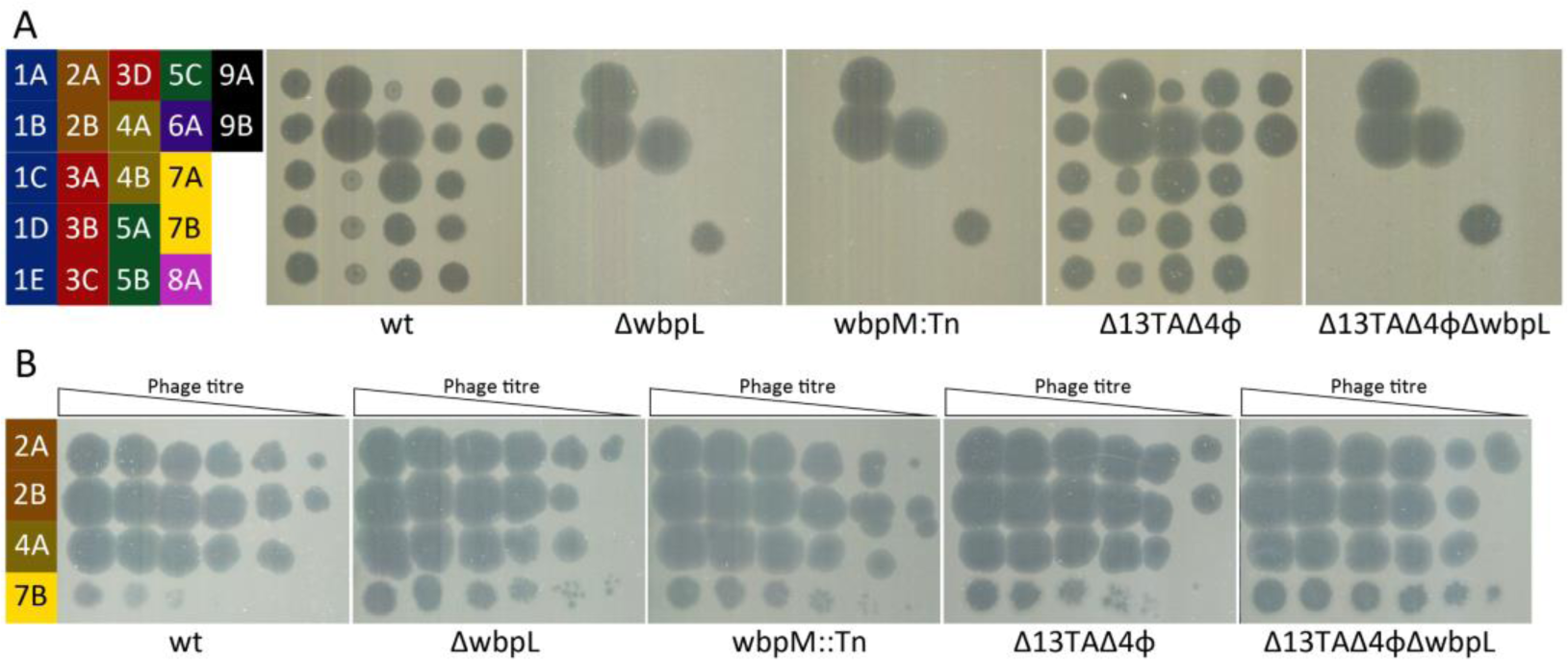
Most CEPEST phages require intact LPS for infection. A: A qualitative assay of phage infection of all 22 phage species representatives on *P. putida* PaW85 wild type (wt) and its *wbpL* or *wbpM* deficient strains and *wbpL* deletion derivative in the Δ13TAΔ4ϕ background. B: EOP of phages that can infect mutants with deficient LPS.

## Discussion

The CEPEST collection of bacteriophages isolated during this work represents the first broad selection of *P. putida* environmental phages. Genome homology analysis allocated the CEPEST collection phages into nine genus clusters with 22 species. As only 13 complete *P. putida* phage genome sequences are available to date (February 2024, NCBI database), it is not surprising that many of the phages are entirely new and even constitute several new genus clusters. From that it seems that extensive studies are still needed to understand the bacteriophage diversity and any new host bacterium used in isolation could result in identifying many new genera of phages.

### The number of phage species is variable in the genus clusters of the CEPEST collection

The composition of the CEPEST collection indicates that it has a high representation of some phage species but is apparently underrepresented in others. Currently, the collection contains phages from only nine genus clusters, and the species abundance in different clusters is highly variable. While the genus cluster G8 is represented by only one species, there are also clusters having over ten isolates defined as the same phage species (identity >95%). For instance, 2A and 5B species clusters are the most overrepresented, containing 17 and 21 phage isolates, respectively. Repeated isolation of 2A phages is not so surprising as G2 phages produce big plaques (Table S3, Supplementary File 2). As plaque sizes positively correlate with lysis rates (Ameh *et al*., 2020), the propagation of these phages could be favoured during the isolation process that involved an enrichment step. However, G4 phages that also form big plaques were isolated from fewer environmental samples. Therefore, phages of cluster G2 are probably more widespread in the different sampling environments than G4 ones. The repeated isolation of genus cluster G5 phages with average-size plaques is more surprising. As our isolation experiments show the greatest abundance of these phages, they seem to be the most frequent culturable *P. putida* phages in environmental samples. Still, to rule out that the enrichment phase of the isolation method was the cause of the recurrent appearance of certain phages, we also carried out phage isolation without the enrichment step by directly plating phages precipitated from environmental samples. The G5 phages were still the most frequently detected in the samples, as was determined by G5-specific PCR analysis (data not shown). From that, we conclude that the G5 is indeed a prevalent cluster of *P. putida* phages in the environment. Interestingly, the G5 phages represent an entirely new genus cluster (Figure 1A). We hypothesize that the reasons for them not having been isolated previously are their very narrow host range (Figure 6A and D) and a temperature-sensitive infection efficiency (Figure 4B), which means that they would have been missed when using different *Pseudomonas* hosts or higher than 25 °C growth temperature for the isolation.

All phages in the current CEPEST collection are tailed double-stranded DNA phages from the order *Caudovirales*. Over three quarters of all the 67 phages have podovirus morphology, and two genus clusters (G1 and G3) are myoviruses. Surprisingly, we did not isolate any siphoviruses, although the morphology is common for *E. coli* phages (Korf *et al*., 2019; Olsen *et al*., 2020; Maffei *et al*., 2021; Nicolas *et al*., 2023). Moreover, the representation of morphotypes of *P. putida* phages in the current collection remarkably differs from that in several coliphage collections: around 70% of lytic coliphages are myoviruses, 22-25% siphoviruses and less than 10% podoviruses (Korf *et al*., 2019; Olsen *et al*., 2020). Interestingly, if we surveyed the distribution of virion morphotypes of previously isolated *P. putida* phages, only myo- and podoviruses were found in accordance with our results. Out of 13 sequenced phages that have been isolated with different *P. putida* strains, four had no morphology data, but six were podoviruses (Lee and Boezi, 1966; Glukhov *et al*., 2012; Ngiam, Weynberg and Guo, 2022) and three myoviruses (Magill *et al*., 2017; Jaryenneh, Schoeniger and Mageeney, 2023). Thus, it seems that *P. putida* phages inherently differ from the phages infecting *E. coli* with podovirus morphotype being much more prevalent.

### The CEPEST collection phages are highly dependent on infection temperature

Most CEPEST collection phages seem to be adapted to low temperatures, as their optimum infection temperature was 15 to 20 °C, and many failed to lyse *P. putida* at higher temperatures (Figure 4). This is actually not surprising, given that *P. putida* is an environmental bacterium that usually meets its natural phage enemies in soil and water at much lower temperatures than 30 °C that is routinely used in the laboratory for growing *P. putida*. Previous studies have shown that temperature may drastically influence the efficiency of phage infection. For instance, coliphages have been divided into three groups according to their temperature sensitivity: high-temperature (HT) phages that lose infectivity at or below 25 °C, low-temperature (LT) phages that lose infectivity at or above 30 °C and mid-temperature (MT) phages that are infectious in the range 15 to 42 °C (Seeley and Primrose, 1980). LT coliphages seem less abundant than HT and MT coliphages (Seeley and Primrose, 1980). However, as the isolation of *E. coli* phages is mostly carried out at 37 °C (Jurczak-Kurek *et al*., 2016; Korf *et al*., 2019; Olsen *et al*., 2020; Maffei *et al*., 2021; Nicolas *et al*., 2023), the LT coliphages might be missing or underrepresented in coliphage collections due to the isolation procedures. This is most probably also the case with the 13 *P. putida* phages isolated so far – as previous isolations were done at 30 °C (Ngiam, Weynberg and Guo, 2022; Jaryenneh, Schoeniger and Mageeney, 2023) or even at 33 °C (Lee and Boezi, 1966), the phages that require lower temperatures might not have been detected. Considering the temperature-sensitivity pattern of the CEPEST collection phages, the only phages that we could theoretically catch at 30 °C would be those from the genus clusters G1, G4, and G8.

Therefore, we would have missed other six genus clusters, among them also the most prevalent ones: clusters G2 and G5. This illustrates that temperature should always be considered when isolating and studying phages of environmental microbes. It has been shown that the temperature profile of phages depends on their origin: phages originating from warm-blooded animals tend to be MT or HT type, while LT-type phages can be found in aquatic habitats (Lee and Boezi, 1966). As our environmental samples originate from the temperate climate of Estonia, it is not surprising that many CEPEST phages displayed an even higher infection efficiency at 15 °C compared to the isolation temperature of 20 °C (Figure 4). Therefore, when expanding the collection, one might consider lowering the isolation temperature further down to 15 °C. The evident low-temperature requirement of many *P. putida* phages is very encouraging when considering *P. putida* as a promising microbial cell factory for biocatalysis. It seems that besides its versatile metabolism and high stress tolerance that makes *P. putida* a valuable candidate for bioproduction, its increased phage resistance at higher temperatures when metabolic reactions are more efficient adds value to its use in bio-industrial applications.

### The host and receptor specificity of the CEPEST collection phages

The majority of the CEPEST collection phages display a narrow host range and could not infect other *Pseudomonas* species or other *P. putida* strains, except for the *P. putida* PaW85 ancestor strain mt-2 (Figure 6). Still, some phages, particularly those from *Pifdecavirus* (G2) and *Ghunavirus* (G4, G7, G9) genus clusters, had a broader host range, as they could lyse several other *P. putida* and even *P. syringae* strains. Phages from new genus clusters G1, G3, and G5 were highly specific to the isolation host, and only one G3 jumbo phage named Peetri could lyse the *P. syringae* B728a strain. Given that the host specificity is very often determined by the level of receptor recognition, it is interesting to note that the only phages able to infect the rough cells of the ΔwbpL strain were from the same genus clusters with a broader host range. Phages from genus cluster G2 had a wide host range and could also infect the O-antigen deficient strain ΔwbpL (Figure 6 and 7). However, opposite to G2 cluster, the phages from genus clusters G4 and G7 that could infect the O-antigen deficient strain, i.e., Emajogi (4A) and Kompost-2 (7B), had a narrow host range, whereas the other species in these clusters, Luke-2 (4B) and Kallioja (7A) with a wider host range, were unable to infect the ΔwbpL strain (Figure 6 and 7). This suggests that the receptors of the phages able to infect the ΔwbpL cells are different. The infection pattern of *Ghunavirus* Kompost-2 is unique in the collection, as the infection is more efficient when the O-antigen is missing. This suggests that the O-antigen probably masks the main receptor of the phage on the wild-type cells, and the receptor is more accessible upon O-antigen deletion.

Previous studies of *Pseudomonas* phage gh-1 from the *Ghunavirus* genus have shown that the infection by this phage is dependent on the O-antigen of bacterial LPS (Kovalyova and Kropinski, 2003). Three genus clusters, G4, G7, and G9, in the CEPEST collection could also be grouped with the *Ghunavirus* (Figure 2A). The closest to *Pseudomonas* phage gh-1 was the genus cluster G4 (overall genome sequence identity around 89%; Table S7, Supplementary File 2). Interestingly, although the sequence identity percentage of the two G4 species to gh-1 is similar, only the 4B phage Luke-2, but not the 4A phage Emajogi, was dependent on the O-antigen for infection (Figure 7). As C-terminal parts of tail fiber proteins are responsible for receptor recognition, it is quite common for these areas of the proteins to undergo shuffling as one of the processes that diversify the host range of the phages (Smug *et al*., 2023). This can lead to otherwise similar phages having diverse receptors or varied host range. The CEPEST collection illustrates this well, as both the host range and the receptor specificity differ between species 4A and 4B (Figure 6 and 7), although their full genome sequences are almost 93% identical (Table S7, Supplementary File 2).

### *P. putida* genomic prophages and plasmid pWW0 confer defence against certain CEPEST collection phages

It is evident that the CEPEST collection contains only phages that can overcome the putative antiphage systems of *P. putida*. The only exceptions are the potential defence provided by the TA systems and cryptic prophages that were deleted from the genome of the isolation host strain (Figure 1). Our previous efforts have not detected any importance of *P. putida* chromosomal TA systems for bacterial physiology (Rosendahl *et al*., 2020). However, due to the lack of *P. putida* phages, we could not previously test the possible effect of TA systems in phage defence. Here, we compared the phage resistance of wild-type *P. putida* PaW85 and its 13 TA system deletion derivative but did not detect any differences in the phage defence of the two strains (Figure 5 A and B). Our previous data has revealed that several TA systems encode for non-toxic proteins (Rosendahl *et al*., 2020), which indicates that these toxins have probably lost their function even if they once would have participated in phage defence. Nevertheless, eight out of the 13 toxins deleted from *P. putida* were moderately toxic or lethal proteins (Rosendahl *et al*., 2020) and still did not affect the phage resistance of the bacterium. This means that none of the phages in the current CEPEST collection can activate any of the 13 TA systems in *P. putida*. However, the 22 species and 67 isolates of phages actually represent a very small sample size of all possible phages. Thus, from these results, we cannot rule out the possibility that some *P. putida* TA systems could be active against yet unknown phages. As the collection is upgraded, the TA systems’ involvement in phage infection will be kept under consideration. Yet, current results demonstrate that the 13 chromosomal TA systems provide no general phage defence and neither do they act as specific phage defence systems against the 67 tested bacteriophage isolates of the CEPEST collection.

Differently from TA systems, however, the four cryptic prophages can protect *P. putida* at variable strength against about half of the 22 species of phages. Our future work will determine whether this protection is provided by a single prophage or several prophages together. It is known that prophages may inhibit the expression of lytic genes from similar phages (Susskind, Wright and Botstein, 1974). As prophages keep their genes for lytic development repressed, the infection of phages that share similar repressors is also inhibited (Bondy-Denomy *et al*., 2016). Yet, as our analysis of the prophage regulator or any other prophage genes revealed no resemblance among any of the phages in the CEPEST collection (data not shown), this type of defence does not seem to be the mechanism of protection by these prophages. Aside from suppressing the lytic development of similar phages, prophages can carry various defence genes that protect the host from attacks by other phages. For example, the superinfection exclusion (Sie) systems either inhibit the adsorption of new phages (Bondy-Denomy *et al*., 2016) or prevent their genome injection (Susskind, Wright and Botstein, 1974; Hofer, Ruge and Dreiseikelmann, 1995; Cumby *et al*., 2012). Prophages can also encode TA gene(s) or other Abi systems (Owen *et al*., 2021; Zhang *et al*., 2022) that prevent the spread of the exogenous phages in bacterial population by inducing the death of the first infected cell. Our next efforts will be directed towards identifying these systems to pinpoint the mechanisms of the prophage-provided defence.

The studies of host specificity of the CEPEST collection phages led to the discovery of the protective effect of the TOL plasmid pWW0 (Worsey and Williams, 1975; Assinder and Williams, 1990; Ramos, Marqués and Timmis, 1997; Greated *et al*., 2002) against the phages from genus cluster G5 (Figure 6C). It has been shown that plasmids may influence the efficiency of phage infection by either plasmid-encoded factors that are required for phage entry (Olsen, Siak and Gray, 1974; Haase *et al*., 1995) or, on the contrary, strongly decreasing the infection efficiency by plasmid-coded defence systems (Emond *et al*., 1998). The pWW0 plasmid-encoded pili have been shown to support the infection of plasmid-dependent phages PR4 and PRD1 (Bradley and Williams, 1982), but to our knowledge, the pWW0 plasmid has never been associated with phage defence. Given, that plasmids have been shown to code for phage defence elements, like abortive infection systems (Bouchard *et al*., 2002) or CRISPR-Cas systems (Pinilla-Redondo *et al*., 2022), it is possible that the pWW0 plasmid contains a phage defence element that protects against phages from G5 genus cluster. As another option, the presence of the TOL plasmid in the cell could somehow change the phage resistance indirectly. The exact mechanisms for the protection provided by the TOL plasmid are subjects of further research using the diverse and well-represented genus cluster G5 (Table S3, Supplementary File 2) of the CEPEST collection.

## Conclusion

Here, we present the first broad collection of phages infecting the environmental bacterium and model organism *P. putida* PaW85. The collection is built up of phages isolated against a host whose phage resistance is completely unknown. The results obtained on the phage defence during the screening stages, will be followed up with research behind the mechanisms of the observed biological effects. Thus, the CEPEST collection allows the phage resistance research of this widely used model bacterium. As knowledge on the phage defence of *P. putida* broadens, more phages will be isolated using weakened *P. putida* strains as hosts to increase the diversity of the CEPEST collection and improve the understanding of the interactions between *P. putida* PaW85 and its phages.

## Acknowledgements

We are grateful to Esteban Martínez-García for kindly providing plasmids for prophage deletion. We are also thankful to Kaida Koppel and Raivo Raid for providing the transmission electron microscopy service, Sander Blei and Anita Lipu for help in phage sampling, and Tanel Ilmjärv and Ingrem Popazova for providing the strain *P. putida* PaW85 [pWW0]. We thank Andres Ainelo for critically reading the manuscript and scientific illustrator Ali Haririan for input to the graphical abstract. We acknowledge the CELMS collection of the Institute of Molecular and Cell Biology, University of Tartu for the environmental bacterial strains.

This work was supported by the Estonian Research Council (grant PRG1431 to RH and grants MOBTP1017 and together with EMBO the EMBO IG-5323-2023 to HT) and ERC (StG PhaBacArms grant no 101116205 to HT).

## References

Ameh, E.M. et al. (2020) ‘Lysis Performance of Bacteriophages with Different Plaque Sizes and Comparison of Lysis Kinetics After Simultaneous and Sequential Phage Addition’, PHAGE, 1(3), pp. 149–157. Available at: 10.1089/phage.2020.0005.

Aparicio, T., De Lorenzo, V. and Martínez-García, E. (2019) ‘Improved Thermotolerance of Genome-Reduced *Pseudomonas putida* EM42 Enables Effective Functioning of the P _L_ / *c* I857 System’, Biotechnology Journal, 14(1), p. 1800483. Available at: 10.1002/biot.201800483.

Assinder, S.J. and Williams, P.A. (1990) ‘The TOL Plasmids: Determinants of the Catabolism of Toluene and the Xylenes’, in Advances in Microbial Physiology. Elsevier, pp. 1–69. Available at: 10.1016/S0065-2911(08)60119-8.

Bayley, S.A. et al. (1977) ‘Two modes of loss of the tol function from Pseudomonas putida mt-2’, Molecular and General Genetics MGG, 154(2), pp. 203–204. Available at: 10.1007/BF00330838.

Bélanger, M., Burrows, L.L. and Lam, J.S. (1999) ‘Functional analysis of genes responsible for the synthesis of the B-band O antigen of Pseudomonas aeruginosa serotype O6 lipopolysaccharide The GenBank accession number for the sequence reported in this paper is AF035937.’, Microbiology, 145(12), pp. 3505–3521. Available at: 10.1099/00221287-145-12-3505.

Bertozzi Silva, J., Storms, Z. and Sauvageau, D. (2016) ‘Host receptors for bacteriophage adsorption’, FEMS Microbiology Letters. Edited by A. Millard, 363(4), p. fnw002. Available at: 10.1093/femsle/fnw002.

Bin Jang, H., et al. (2019) ‘Taxonomic assignment of uncultivated prokaryotic virus genomes is enabled by gene-sharing networks’, Nature Biotechnology, 37(6), pp. 632–639. Available at: 10.1038/s41587-019-0100-8.

Bogosian, G. (2005) ‘Control of Bacteriophage in Commercial Microbiology and Fermentation Facilities’, in R. Calendar and S.T. Abedon (eds) The Bacteriophages. Oxford University PressNew York, NY, pp. 667–673. Available at: 10.1093/oso/9780195148503.003.0042.

Bondy-Denomy, J. et al. (2016) ‘Prophages mediate defense against phage infection through diverse mechanisms’, The ISME Journal, 10(12), pp. 2854–2866. Available at: 10.1038/ismej.2016.79.

Bouchard, J.D. et al. (2002) ‘Characterization of the Two-Component Abortive Phage Infection Mechanism AbiT from *Lactococcus lactis*’, Journal of Bacteriology, 184(22), pp. 6325–6332. Available at: 10.1128/JB.184.22.6325-6332.2002.

Bradley, D.E. and Williams, P.A. (1982) ‘The TOL Plasmid is Naturally Derepressed for Transfer’, Microbiology, 128(12), pp. 3019–3024. Available at: 10.1099/00221287-128-12-3019.

Chen, S. et al. (2018) ‘fastp: an ultra-fast all-in-one FASTQ preprocessor’, Bioinformatics, 34(17), pp. i884–i890. Available at: 10.1093/bioinformatics/bty560.

Comeau, A.M. et al. (2008) ‘Exploring the prokaryotic virosphere’, Research in Microbiology, 159(5), pp. 306–313. Available at: 10.1016/j.resmic.2008.05.001.

Cook, R. et al. (2021) ‘INfrastructure for a PHAge REference Database: Identification of Large-Scale Biases in the Current Collection of Cultured Phage Genomes’, PHAGE, 2(4), pp. 214–223. Available at: 10.1089/phage.2021.0007.

Creuzenet, C. and Lam, J.S. (2001) ‘Topological and functional characterization of WbpM, an inner membrane UDP-GlcNAc C _6_ dehydratase essential for lipopolysaccharide biosynthesis in *Pseudomonas aeruginosa*’, Molecular Microbiology, 41(6), pp. 1295–1310. Available at: 10.1046/j.1365-2958.2001.02589.x.

Cumby, N. et al. (2012) ‘The Bacteriophage HK97 gp15 Moron Element Encodes a Novel Superinfection Exclusion Protein’, Journal of Bacteriology, 194(18), pp. 5012–5019. Available at: 10.1128/JB.00843-12.

Darling, A.C.E. et al. (2004) ‘Mauve: Multiple Alignment of Conserved Genomic Sequence With Rearrangements’, Genome Research, 14(7), pp. 1394–1403. Available at: 10.1101/gr.2289704.

Dedrick, R.M. et al. (2017) ‘Prophage-mediated defence against viral attack and viral counter-defence’, Nature Microbiology, 2(3), p. 16251. Available at: 10.1038/nmicrobiol.2016.251.

Doron, S. et al. (2018) ‘Systematic discovery of antiphage defense systems in the microbial pangenome’, Science, 359(6379), p. eaar4120. Available at: 10.1126/science.aar4120.

Dos Santos, V.A.P.M., et al. (2004) ‘Insights into the genomic basis of niche specificity of *Pseudomonas putida* KT2440’, Environmental Microbiology, 6(12), pp. 1264–1286. Available at: 10.1111/j.1462-2920.2004.00734.x.

Dy, R.L. et al. (2014) ‘A widespread bacteriophage abortive infection system functions through a Type IV toxin–antitoxin mechanism’, Nucleic Acids Research, 42(7), pp. 4590–4605. Available at: 10.1093/nar/gkt1419.

Edgar, R.C. (2004) ‘MUSCLE: multiple sequence alignment with high accuracy and high throughput’, Nucleic Acids Research, 32(5), pp. 1792–1797. Available at: 10.1093/nar/gkh340.

Emond, E. et al. (1998) ‘AbiQ, an Abortive Infection Mechanism from *Lactococcus lactis*’, Applied and Environmental Microbiology, 64(12), pp. 4748–4756. Available at: 10.1128/AEM.64.12.4748-4756.1998.

Fineran, P.C. et al. (2009) ‘The phage abortive infection system, ToxIN, functions as a protein–RNA toxin– antitoxin pair’, Proceedings of the National Academy of Sciences, 106(3), pp. 894–899. Available at: 10.1073/pnas.0808832106.

Fraikin, N. and Van Melderen, L. (2024) ‘Single-cell evidence for plasmid addiction mediated by toxin– antitoxin systems’, Nucleic Acids Research, 52(4), pp. 1847–1859. Available at: 10.1093/nar/gkae018.

Garneau, J.R. et al. (2017) ‘PhageTerm: a tool for fast and accurate determination of phage termini and packaging mechanism using next-generation sequencing data’, Scientific Reports, 7(1), p. 8292. Available at: 10.1038/s41598-017-07910-5.

Georjon, H. and Bernheim, A. (2023) ‘The highly diverse antiphage defence systems of bacteria’, Nature Reviews Microbiology, 21(10), pp. 686–700. Available at: 10.1038/s41579-023-00934-x.

Gilchrist, C.L.M. and Chooi, Y.-H. (2021) ‘clinker & clustermap.js: automatic generation of gene cluster comparison figures’, Bioinformatics. Edited by P. Robinson, 37(16), pp. 2473–2475. Available at: 10.1093/bioinformatics/btab007.

Glukhov, A.S., et al. (2012) ‘Genomic Analysis of Pseudomonas putida Phage tf with Localized Single-Strand DNA Interruptions’, PLoS ONE. Edited by A.R. Poteete, 7(12), p. e51163. Available at: 10.1371/journal.pone.0051163.

Greated, A. et al. (2002) ‘Complete sequence of the IncP-9 TOL plasmid pWW0 from *Pseudomonas putida*’, Environmental Microbiology, 4(12), pp. 856–871. Available at: 10.1046/j.1462-2920.2002.00305.x.

Guegler, C.K. and Laub, M.T. (2021) ‘Shutoff of host transcription triggers a toxin-antitoxin system to cleave phage RNA and abort infection’, Molecular Cell, 81(11), pp. 2361–2373.e9. Available at: 10.1016/j.molcel.2021.03.027.

Haase, J. et al. (1995) ‘Bacterial conjugation mediated by plasmid RP4: RSF1010 mobilization, donor-specific phage propagation, and pilus production require the same Tra2 core components of a proposed DNA transport complex’, Journal of Bacteriology, 177(16), pp. 4779–4791. Available at: 10.1128/jb.177.16.4779-4791.1995.

Hazan, R. and Engelberg-Kulka, H. (2004) ‘Escherichia coli mazEF-mediated cell death as a defense mechanism that inhibits the spread of phage P1’, Molecular Genetics and Genomics, 272(2), pp. 227–234. Available at: 10.1007/s00438-004-1048-y.

Hofer, B., Ruge, M. and Dreiseikelmann, B. (1995) ‘The superinfection exclusion gene (sieA) of bacteriophage P22: identification and overexpression of the gene and localization of the gene product’, Journal of Bacteriology, 177(11), pp. 3080–3086. Available at: 10.1128/jb.177.11.3080-3086.1995.

Horvath, P. and Barrangou, R. (2010) ‘CRISPR/Cas, the Immune System of Bacteria and Archaea’, Science, 327(5962), pp. 167–170. Available at: 10.1126/science.1179555.

Jaryenneh, J., Schoeniger, J.S. and Mageeney, C.M. (2023) ‘Genome sequence and characterization of a novel Pseudomonas putida phage, MiCath’, Scientific Reports, 13(1), p. 21834. Available at: 10.1038/s41598-023-48634-z.

Jurczak-Kurek, A. et al. (2016) ‘Biodiversity of bacteriophages: morphological and biological properties of a large group of phages isolated from urban sewage’, Scientific Reports, 6(1), p. 34338. Available at: 10.1038/srep34338.

Kalyaanamoorthy, S. et al. (2017) ‘ModelFinder: fast model selection for accurate phylogenetic estimates’, Nature Methods, 14(6), pp. 587–589. Available at: 10.1038/nmeth.4285.

Kelly, A. et al. (2023) ‘Toxin–antitoxin systems as mediators of phage defence and the implications for abortive infection’, Current Opinion in Microbiology, 73, p. 102293. Available at: 10.1016/j.mib.2023.102293.

Korf, I.H.E. et al. (2019) ‘Still Something to Discover: Novel Insights into Escherichia coli Phage Diversity and Taxonomy’, Viruses, 11(5), p. 454. Available at: 10.3390/v11050454.

Kovalyova, I.V. and Kropinski, A.M. (2003) ‘The complete genomic sequence of lytic bacteriophage gh-1 infecting Pseudomonas putida—evidence for close relationship to the T7 group’, Virology, 311(2), pp. 305–315. Available at: 10.1016/S0042-6822(03)00124-7.

Lee, L.F. and Boezi, J.A. (1966) ‘Characterization of Bacteriophage gh-1 for *Pseudomonas putida*’, Journal of Bacteriology, 92(6), pp. 1821–1827. Available at: 10.1128/jb.92.6.1821-1827.1966.

LeRoux, M. et al. (2022) ‘The DarTG toxin-antitoxin system provides phage defence by ADP-ribosylating viral DNA’, Nature Microbiology, 7(7), pp. 1028–1040. Available at: 10.1038/s41564-022-01153-5.

LeRoux, M. and Laub, M.T. (2022) ‘Toxin-Antitoxin Systems as Phage Defense Elements’, Annual Review of Microbiology, 76(1), pp. 21–43. Available at: 10.1146/annurev-micro-020722-013730.

Letunic, I. and Bork, P. (2021) ‘Interactive Tree Of Life (iTOL) v5: an online tool for phylogenetic tree display and annotation’, Nucleic Acids Research, 49(W1), pp. W293–W296. Available at: 10.1093/nar/gkab301.

Lindberg, A.A. (1973) ‘Bacteriophage Receptors’, Annual Review of Microbiology, 27(1), pp. 205–241. Available at: 10.1146/annurev.mi.27.100173.001225.

Lopatina, A., Tal, N. and Sorek, R. (2020) ‘Abortive Infection: Bacterial Suicide as an Antiviral Immune Strategy’, Annual Review of Virology, 7(1), pp. 371–384. Available at: 10.1146/annurev-virology-011620-040628.

Los, M. (2012) ‘Minimization and Prevention of Phage Infections in Bioprocesses’, in Q. Cheng (ed.) Microbial Metabolic Engineering. New York, NY: Springer New York (Methods in Molecular Biology), pp. 305–315. Available at: 10.1007/978-1-61779-483-4_19.

Maffei, E., et al. (2021) ‘Systematic exploration of Escherichia coli phage–host interactions with the BASEL phage collection’, PLOS Biology. Edited by J. Barr, 19(11), p. e3001424. Available at: 10.1371/journal.pbio.3001424.

Magill, D.J., et al. (2017) ‘Pf16 and phiPMW: Expanding the realm of Pseudomonas putida bacteriophages’, PLOS ONE. Edited by M.J. Van Raaij, 12(9), p. e0184307. Available at: 10.1371/journal.pone.0184307.

Makarova, K.S. et al. (2011) ‘Defense Islands in Bacterial and Archaeal Genomes and Prediction of Novel Defense Systems’, Journal of Bacteriology, 193(21), pp. 6039–6056. Available at: 10.1128/JB.05535-11.

Martínez-García, E. et al. (2014) ‘Pseudomonas 2.0: genetic upgrading of P. putida KT2440 as an enhanced host for heterologous gene expression’, Microbial Cell Factories, 13(1), p. 159. Available at: 10.1186/s12934-014-0159-3.

Martínez-García, E. et al. (2015) ‘Freeing *Pseudomonas putida* KT2440 of its proviral load strengthens endurance to environmental stresses: The prophages of *P. putida* KT2440’, Environmental Microbiology, 17(1), pp. 76–90. Available at: 10.1111/1462-2920.12492.

Martínez-García, E. and De Lorenzo, V. (2024) ‘Pseudomonas putida as a synthetic biology chassis and a metabolic engineering platform’, Current Opinion in Biotechnology, 85, p. 103025. Available at: 10.1016/j.copbio.2023.103025.

Martínez-García, E. and de Lorenzo, V. (2011) ‘Engineering multiple genomic deletions in Gram-negative bacteria: analysis of the multi-resistant antibiotic profile of *Pseudomonas putida* KT2440: Tools for editing Gram-negative genomes’, Environmental Microbiology, 13(10), pp. 2702–2716. Available at: 10.1111/j.1462-2920.2011.02538.x.

Millman, A. et al. (2022) ‘An expanded arsenal of immune systems that protect bacteria from phages’, Cell Host & Microbe, 30(11), pp. 1556–1569.e5. Available at: 10.1016/j.chom.2022.09.017.

Moraru, C., Varsani, A. and Kropinski, A.M. (2020) ‘VIRIDIC—A Novel Tool to Calculate the Intergenomic Similarities of Prokaryote-Infecting Viruses’, Viruses, 12(11), p. 1268. Available at: 10.3390/v12111268.

Ngiam, L., Weynberg, K.D. and Guo, J. (2022) ‘The presence of plasmids in bacterial hosts alters phage isolation and infectivity’, ISME Communications, 2(1), p. 75. Available at: 10.1038/s43705-022-00158-9.

Nguyen, L.-T. et al. (2015) ‘IQ-TREE: A Fast and Effective Stochastic Algorithm for Estimating Maximum-Likelihood Phylogenies’, Molecular Biology and Evolution, 32(1), pp. 268–274. Available at: 10.1093/molbev/msu300.

Nicolas, M., et al. (2023) ‘Isolation and Characterization of a Novel Phage Collection against Avian-Pathogenic *Escherichia coli*’, Microbiology Spectrum. Edited by T.G. Denes, 11(3), pp. e04296-22. Available at: 10.1128/spectrum.04296-22.

Nikel, P.I. et al. (2016) ‘From dirt to industrial applications: *Pseudomonas putida* as a Synthetic Biology chassis for hosting harsh biochemical reactions’, Current Opinion in Chemical Biology, 34, pp. 20–29. Available at: 10.1016/j.cbpa.2016.05.011.

Nobrega, F.L. et al. (2018) ‘Targeting mechanisms of tailed bacteriophages’, Nature Reviews Microbiology, 16(12), pp. 760–773. Available at: 10.1038/s41579-018-0070-8.

Ogura, T. and Hiraga, S. (1983) ‘Mini-F plasmid genes that couple host cell division to plasmid proliferation.’, Proceedings of the National Academy of Sciences, 80(15), pp. 4784–4788. Available at: 10.1073/pnas.80.15.4784.

Olsen, N.S. et al. (2020) ‘Exploring the Remarkable Diversity of Culturable Escherichia coli Phages in the Danish Wastewater Environment’, Viruses, 12(9), p. 986. Available at: 10.3390/v12090986.

Olsen, R.H., Siak, J.-S. and Gray, R.H. (1974) ‘Characteristics of PRD1, a Plasmid-Dependent Broad Host Range DNA Bacteriophage’, Journal of Virology, 14(3), pp. 689–699. Available at: 10.1128/jvi.14.3.689-699.1974.

Owen, S.V. et al. (2021) ‘Prophages encode phage-defense systems with cognate self-immunity’, Cell Host & Microbe, 29(11), pp. 1620–1633.e8. Available at: 10.1016/j.chom.2021.09.002.

Patel, P.H. and Maxwell, K.L. (2023) ‘Prophages provide a rich source of antiphage defense systems’, Current Opinion in Microbiology, 73, p. 102321. Available at: 10.1016/j.mib.2023.102321.

Pinilla-Redondo, R. et al. (2022) ‘CRISPR-Cas systems are widespread accessory elements across bacterial and archaeal plasmids’, Nucleic Acids Research, 50(8), pp. 4315–4328. Available at: 10.1093/nar/gkab859.

Prjibelski, A. et al. (2020) ‘Using SPAdes De Novo Assembler’, Current Protocols in Bioinformatics, 70(1), p. e102. Available at: 10.1002/cpbi.102.

Ramos, J.L., Marqués, S. and Timmis, K.N. (1997) ‘TRANSCRIPTIONAL CONTROL OF THE *PSEUDOMONAS* TOL PLASMID CATABOLIC OPERONS IS ACHIEVED THROUGH AN INTERPLAY OF HOST FACTORS AND PLASMID-ENCODED REGULATORS’, Annual Review of Microbiology, 51(1), pp. 341–373. Available at: 10.1146/annurev.micro.51.1.341.

Regenhardt, D. et al. (2002) ‘Pedigree and taxonomic credentials of *Pseudomonas putida* strain KT2440’, Environmental Microbiology, 4(12), pp. 912–915. Available at: 10.1046/j.1462-2920.2002.00368.x.

Reva, O.N. et al. (2006) ‘Functional Genomics of Stress Response in *Pseudomonas putida* KT2440’, Journal of Bacteriology, 188(11), pp. 4079–4092. Available at: 10.1128/JB.00101-06.

Rocchetta, H.L. et al. (1998) ‘Three rhamnosyltransferases responsible for assembly of the A-band D - rhamnan polysaccharide in *Pseudomonas aeruginosa* : a fourth transferase, WbpL, is required for the initiation of both A-band and B-band lipopolysaccharide synthesis’, Molecular Microbiology, 28(6), pp. 1103–1119. Available at: 10.1046/j.1365-2958.1998.00871.x.

Rosendahl, S. et al. (2020) ‘Chromosomal toxin-antitoxin systems in *Pseudomonas putida* are rather selfish than beneficial’, Scientific Reports, 10(1), p. 9230. Available at: 10.1038/s41598-020-65504-0.

Seeley, N.D. and Primrose, S.B. (1980) ‘The Effect of Temperature on the Ecology of Aquatic Bacteriophages’, Journal of General Virology, 46(1), pp. 87–95. Available at: 10.1099/0022-1317-46-1-87.

Seemann, T. (2014) ‘Prokka: rapid prokaryotic genome annotation’, Bioinformatics, 30(14), pp. 2068–2069. Available at: 10.1093/bioinformatics/btu153.

Sharma, R.C. and Schimke, R.T. (1996) ‘Preparation of Electro-Competent *E. coli* Using Salt-Free Growth Medium’, BioTechniques, 20(1), pp. 42–44. Available at: 10.2144/96201bm08.

Short, F.L. et al. (2018) ‘The bacterial Type III toxin-antitoxin system, ToxIN, is a dynamic protein-RNA complex with stability-dependent antiviral abortive infection activity’, Scientific Reports, 8(1), p. 1013. Available at: 10.1038/s41598-017-18696-x.

Smug, B.J. et al. (2023) ‘Ongoing shuffling of protein fragments diversifies core viral functions linked to interactions with bacterial hosts’, Nature Communications, 14(1), p. 7460. Available at: 10.1038/s41467-023-43236-9.

Song, S. and Wood, T.K. (2020) ‘A Primary Physiological Role of Toxin/Antitoxin Systems Is Phage Inhibition’, Frontiers in Microbiology, 11, p. 1895. Available at: 10.3389/fmicb.2020.01895.

Susskind, M.M., Wright, A. and Botstein, D. (1974) ‘Superinfection exclusion by P22 prophage in lysogens of Salmonella typhimurium’, Virology, 62(2), pp. 367–384. Available at: 10.1016/0042-6822(74)90399-7.

Terzian, P. et al. (2021) ‘PHROG: families of prokaryotic virus proteins clustered using remote homology’, NAR Genomics and Bioinformatics, 3(3), p. lqab067. Available at: 10.1093/nargab/lqab067.

Tock, M.R. and Dryden, D.T. (2005) ‘The biology of restriction and anti-restriction’, Current Opinion in Microbiology, 8(4), pp. 466–472. Available at: 10.1016/j.mib.2005.06.003.

Vassallo, C.N. et al. (2022) ‘A functional selection reveals previously undetected anti-phage defence systems in the E. coli pangenome’, Nature Microbiology, 7(10), pp. 1568–1579. Available at: 10.1038/s41564-022-01219-4.

Volke, D.C. et al. (2020) ‘Synthetic control of plasmid replication enables target- and self-curing of vectors and expedites genome engineering of Pseudomonas putida’, Metabolic Engineering Communications, 10, p. e00126. Available at: 10.1016/j.mec.2020.e00126.

Worsey, M.J. and Williams, P.A. (1975) ‘Metabolism of toluene and xylenes by Pseudomonas (putida (arvilla) mt-2: evidence for a new function of the TOL plasmid’, Journal of Bacteriology, 124(1), pp. 7–13. Available at: 10.1128/jb.124.1.7-13.1975.

Wozniak, R.A.F. and Waldor, M.K. (2009) ‘A Toxin–Antitoxin System Promotes the Maintenance of an Integrative Conjugative Element’, PLoS Genetics. Edited by J. Casadesús, 5(3), p. e1000439. Available at: 10.1371/journal.pgen.1000439.

Zhang, T. et al. (2022) ‘Direct activation of a bacterial innate immune system by a viral capsid protein’, Nature [Preprint]. Available at: 10.1038/s41586-022-05444-z.

