## Supplementary Figures S1-S3 for "Isolation and characterization of a phage collection against *Pseudomonas putida*"

**Supplementary File 3**

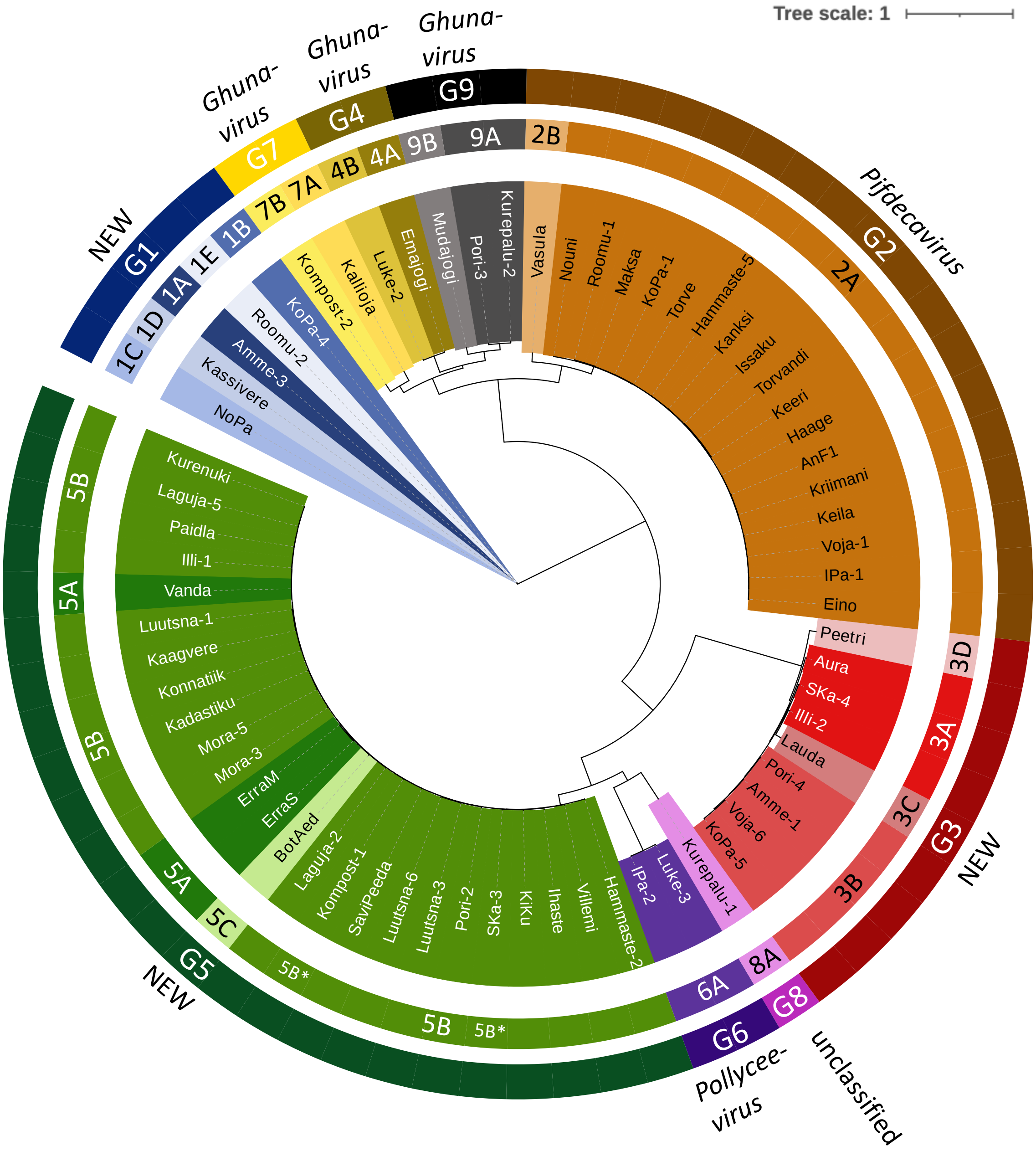


**Figure S1. Phylogenetic tree of all 67 isolates in CEPEST library.** Tree is based on Muscle alignment of four concatenated protein sequences (major head protein, DNA primase, spanin and terminase large subunit), calculated with IQ-TREE using ModelFinder and visualized using iTOL online tool.

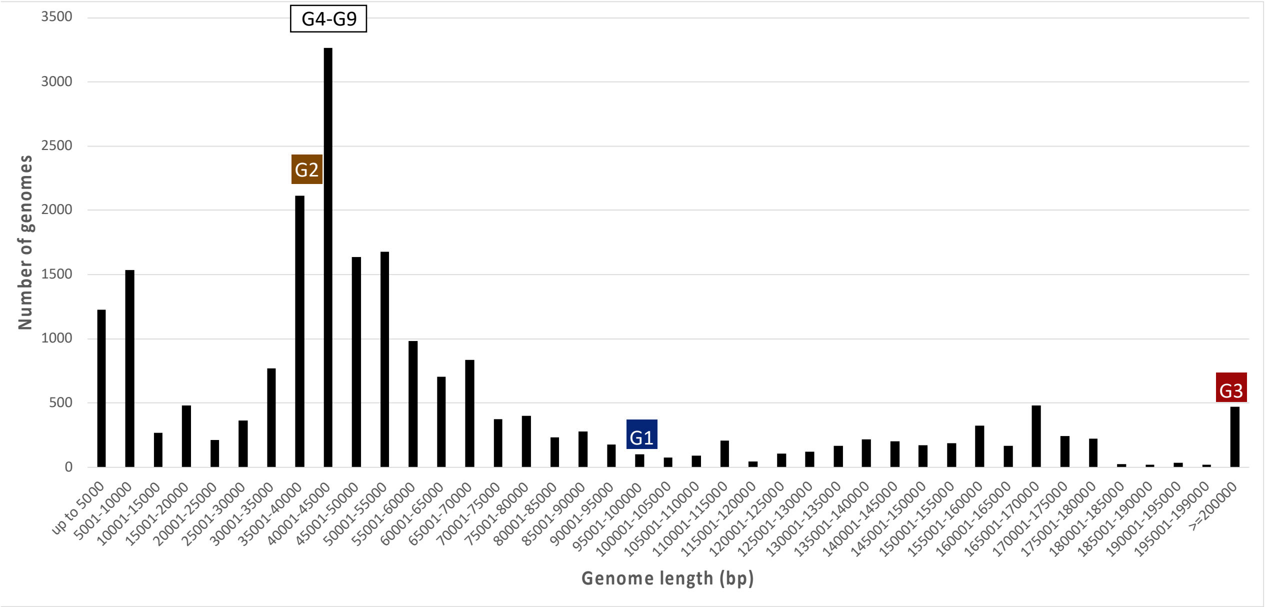


**Figure S2.** CEPEST phages genome length compared to genome length distribution of published complete phage genomes (based on Inphared database collection, February 1st 2024).

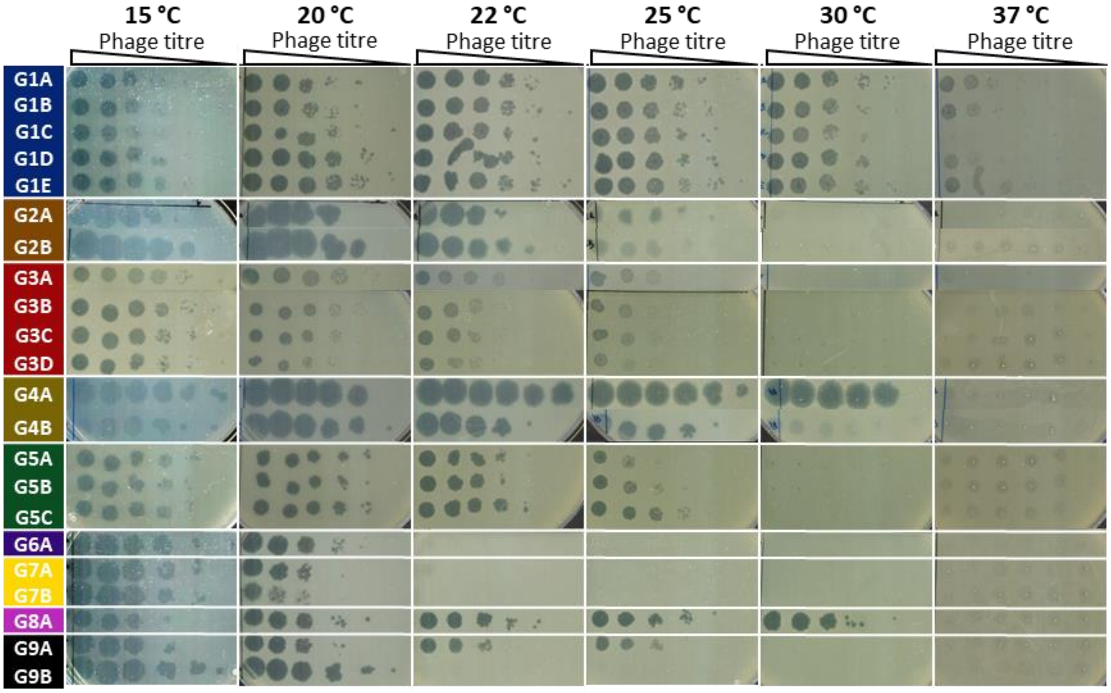


**Figure S3. Temperature sensitivity of all species of CEPEST phages.** The infection of all representative phages was tested at temperatuures ranging from 15 to 37 °C. The phage solutions with $10^8$ titre were serially diluted. And spotted on the plate as 1.5 µL drops. Plaques were recorded after incubation at indicated temperatuure after 20 h (for G3 at 15 °C after 44 h), but monitored for 44 h. The picture is a representative of three independent experiments.
