## Supplementary Table S1 and S2 for "Isolation and characterization of a phage collection against *Pseudomonas putida*"

**Supplementary File 1.**

**Table S1.** Strains and plasmids

| Strain or plasmid | Genotype or characteristic(s) | Source or reference |
|---|---|---|
| ***E. coli* strain** | | |
| DH5α λ*pir* | λpir lysogen of DH5α | (Martínez-García and de Lorenzo, 2011) |
| ***P. putida* strains** | | |
| PaW85 | Wild-type, isogenic to KT2440 | (Bayley *et al.*, 1977) |
| Δ13TA | PaW85 devoid of 13 toxin-antitoxin loci | (Rosendahl *et al.*, 2020) |
| ΔP1 | PaW85 with spontaneous deletion of prophage P1 | this study |
| Δ13TAΔP1 | Δ13TA with spontaneous deletion of prophage P1 | this study |
| ΔP1ΔP4 | ΔP1 with deletion of prophage P4 | this study |
| Δ13TAΔP1ΔP4 | Δ13TAΔP1 with deletion of prophage P4 | this study |
| ΔP1ΔP4ΔP3 | ΔP1ΔP4 with deletion of prophage P3 | this study |
| Δ13TAΔP1ΔP4P3 | Δ13TAΔP1ΔP4 with deletion of prophage P3 | this study |
| Δ4ϕ | ΔP1ΔP4ΔP3 with deletion of prophage P2 | this study |
| Δ13TAΔ4ϕ | Δ13TAΔP1ΔP4P3 with deletion of prophage P2 | this study |
| ΔwbpL | PaW85 with deleted *wbpL* (PP_1804) | this study |
| Δ13TAΔ4ϕΔwbpL | Δ13TAΔ4ϕ with deleted *wbpL* (PP_1804) | this study |
| wbpM::Tn | *wbpM* (PP_1805) in PaW85 is disrupted by Tn5SmlacItac minitransposon ($Sm^r$) | lab collection |
| mt-2 | PaW85 ancestor strain harbouring the pWW0 plasmid | (Worsey and Williams, 1975) |
| PaW85[pWW0] | PaW85 with pWW0 plasmid | lab collection |
| CRTN6 | CELMS collection, isolated from surface seawater | (Viggor *et al.*, 2015) |
| PC13 | CELMS collection, isolated from freshwater | (Heinaru *et al.*, 2000) |
| T9 | CELMS collection, isolated from artificial pond | Merike Jõesaar |
| G7 | CELMS collection, isolated from soil | (Stanier, Palleroni and Doudoroff, 1966) |
| KP49 | CELMS collection, isolated from outflow of wastewater treatment plant | Signe Viggor |
| P63a | CELMS collection, isolated from phenol-polluted river water | Eeva Heinaru |
| Sal3V | CELMS collection, isolated from subsurface water | Eeva Heinaru |
| 1S7 | CELMS collection, isolated from river water | Merike Jõesaar |
| A8 | CELMS collection, isolated from surface seawater | (Jutkina *et al.*, 2011) |
| VAS1 | CELMS collection, isolated from river water | Merike Jõesaar |
| KAR36 | CELMS collection, isolated from potato var. Ants tuber | (Kõiv *et al.*, 2015) |
| ***P. syringae* strains** | | |
| P82 | CELMS collection, isolated from phenol-polluted river water | Eeva Heinaru |
| pv tomato DC3000 | CELMS collection, model plant pathogen, endophyte | (Cuppels, 1986) |
| B728a | CELMS collection, isolated from snap bean leaflet, epiphyte | (Feil *et al.*, 2005) |
| Asal1T | CELMS collection, isolated from surface seawater | Eeva Heinaru |
| DEG | CELMS collection, isolated from subsurface water | Eeva Heinaru |

| | | |
|---|---|---|
| KM2R4 | CELMS collection, isolated from birch sap | Eeva Heinaru |
| RS3 | CELMS collection, isolated from an overgrown pond by the railway | Merike Jõesaar |
| KAR24 | CELMS collection, isolated from potato var. Ants tuber | (Kõiv *et al.*, 2015) |
| ***P. fluorescens* strains** | | |
| Pf0-1 | CELMS collection, isolated from soil | (Compeau *et al.*, 1988) |
| KAR29 | CELMS collection, isolated from potato var. Ants tuber | (Kõiv *et al.*, 2015) |
| ***P. aeruginosa* strains** | | |
| PAO-1L | PAO1 subline, University of Lausanne, Dieter Haas collection | Stephan Heeb |
| KAR21 | CELMS collection, isolated from potato var. Ants tuber | (Kõiv *et al.*, 2015) |
| ***P. stutzeri* strains** | | |
| 37EPH | CELMS collection, isolated from soil | Eeva Heinaru |
| 2C23 | CELMS collection, isolated from surface seawater | (Vedler *et al.*, 2013) |
| **Plasmids** | | |
| pEMG | Plasmid for homologous recombination, *lacZα* with two flanking I-SceI sites ($Km^r$) | (Martínez-García and de Lorenzo, 2011) |
| pSW(I-SceI) | Plasmid coding for I-SceI endonuclease for allelic exchange experiments ($Bp^r$) | (Wong and Mekalanos, 2000) |
| pEMG-P4 | Plasmid containing chimeric DNA fragment for deleting prophage P4 ($Km^r$) | (Martínez-García *et al.*, 2015) |
| pEMG-R3 | Plasmid containing chimeric DNA fragment for deleting prophage P3 ($Km^r$) | (Martínez-García *et al.*, 2015) |
| pEMG-R2 | Plasmid containing chimeric DNA fragment for deleting prophage P2 ($Km^r$) | (Martínez-García *et al.*, 2015) |
| pSNW2 | pEMG derivate with $P_{14g}$*(BCD2)→msfGFP* ($Km^r$) | (Volke *et al.*, 2020) |
| pSNW2-ΔwbpL | pSNW2 containing chimeric DNA fragment for deleting *wbpL* ($Km^r$) | this study |
| pEMG-ΔhicAB2 | pEMG containing chimeric DNA fragment for deleting *hicAB2* ($Km^r$) | this study |

**Table S2.** Oligonucleotides

| Name | Sequence (5'-3')[a] | Use |
|---|---|---|
| 3899Eco | cggaattctacatagccgccctgg | construction of pEMG-ΔhicAB2 |
| 3899TAdel-pikk | ctgtcatagcatcgcaactagggaagcggatgagcct | construction of pEMG-ΔhicAB2 |
| 3900start | tagttgcgatgctatgacag | construction of pEMG-ΔhicAB2 |
| 3901Bam | gtggatccggtgatggagcgagt | construction of pEMG-ΔhicAB2 |
| del1804Eco | aagaattcatggaagtggtggtgatac | construction of pSNW2-ΔwbpL |
| del1804 | cagcacaatcagccatatca | construction of pSNW2-ΔwbpL |
| del1804-pikk | tgatatggctgattgtgctgtaggatggccaagcacgcaa | construction of pSNW2-ΔwbpL |
| del1804Bam | aaggatccaccaatgatgaccatgct | construction of pSNW2-ΔwbpL |
| pp3848 | acaccagccagcacctttc | verification of prophage P1 deletion |
| 3922-r | atcgtcagtgccccattcg | verification of prophage P1 deletion |
| 3025lopp | cgtgtggcgagcaatatcg | verification of prophage P2 deletion |
| 3067sees | gcagctttcgcttgaagttc | verification of prophage P2 deletion |
| PP_t39 | ggacgtggtgaaattggtag | verification of prophage P3 deletion |
| 2298alg | tgcaatcatcgtggtgtcct | verification of prophage P3 deletion |
| graTees | atgtgaccggagcttgcga | verification of prophage P4 deletion |
| 1530R2 | taaccgagaacaggggctac | verification of prophage P4 deletion |

[a] The sites of restriction enzymes used in cloning are underlined.

### References

Bayley, S.A. *et al.* (1977) ‘Two modes of loss of the tol function from Pseudomonas putida mt-2’, *Molecular and General Genetics MGG*, 154(2), pp. 203–204. Available at: https://doi.org/10.1007/BF00330838.

Compeau, G. *et al.* (1988) ‘Survival of rifampin-resistant mutants of Pseudomonas fluorescens and Pseudomonas putida in soil systems’, *Applied and Environmental Microbiology*, 54(10), pp. 2432–2438. Available at: https://doi.org/10.1128/aem.54.10.2432-2438.1988.

Cuppels, D.A. (1986) ‘Generation and Characterization of Tn *5* Insertion Mutations in *Pseudomonas syringae* pv. *tomato*’, *Applied and Environmental Microbiology*, 51(2), pp. 323–327. Available at: https://doi.org/10.1128/aem.51.2.323-327.1986.

Feil, H. *et al.* (2005) ‘Comparison of the complete genome sequences of *Pseudomonas syringae* pv. *syringae* B728a and pv. *tomato* DC3000’, *Proceedings of the National Academy of Sciences*, 102(31), pp. 11064–11069. Available at: https://doi.org/10.1073/pnas.0504930102.

Heinaru, E. *et al.* (2000) ‘Three types of phenol and p-cresol catabolism in phenol- and p-cresol-degrading bacteria isolated from river water continuously polluted with phenolic compounds’, *FEMS Microbiology Ecology*, 31(3), pp. 195–205. Available at: https://doi.org/10.1111/j.1574-6941.2000.tb00684.x.

Jutkina, J. *et al.* (2011) ‘Occurrence of Plasmids in the Aromatic Degrading Bacterioplankton of the Baltic Sea’, *Genes*, 2(4), pp. 853–868. Available at: https://doi.org/10.3390/genes2040853.

Kõiv, V. *et al.* (2015) ‘Microbial population dynamics in response to Pectobacterium atrosepticum infection in potato tubers’, *Scientific Reports*, 5(1), p. 11606. Available at: https://doi.org/10.1038/srep11606.

Martínez-García, E. *et al.* (2015) ‘Freeing *Pseudomonas putida* KT2440 of its proviral load strengthens endurance to environmental stresses: The prophages of *P. putida* KT2440’, *Environmental Microbiology*, 17(1), pp. 76–90. Available at: https://doi.org/10.1111/1462-2920.12492.

Martínez-García, E. and de Lorenzo, V. (2011) ‘Engineering multiple genomic deletions in Gram-negative bacteria: analysis of the multi-resistant antibiotic profile of *Pseudomonas putida* KT2440: Tools for editing Gram-negative genomes’, *Environmental Microbiology*, 13(10), pp. 2702–2716. Available at: https://doi.org/10.1111/j.1462-2920.2011.02538.x.

Rosendahl, S. *et al.* (2020) ‘Chromosomal toxin-antitoxin systems in *Pseudomonas putida* are rather selfish than beneficial’, *Scientific Reports*, 10(1), p. 9230. Available at: https://doi.org/10.1038/s41598-020-65504-0.

Stanier, R.Y., Palleroni, N.J. and Doudoroff, M. (1966) ‘The Aerobic Pseudomonads a Taxonomic Study’, *Journal of General Microbiology*, 43(2), pp. 159–271. Available at: https://doi.org/10.1099/00221287-43-2-159.

Vedler, E. *et al.* (2013) ‘Limnobacter spp. as newly detected phenol-degraders among Baltic Sea surface water bacteria characterised by comparative analysis of catabolic genes’, *Systematic and Applied Microbiology*, 36(8), pp. 525–532. Available at: https://doi.org/10.1016/j.syapm.2013.07.004.

Viggor, S. *et al.* (2015) ‘Occurrence of diverse alkane hydroxylase alkB genes in indigenous oil-degrading bacteria of Baltic Sea surface water’, *Marine Pollution Bulletin*, 101(2), pp. 507–516. Available at: https://doi.org/10.1016/j.marpolbul.2015.10.064.

Volke, D.C. *et al.* (2020) ‘Synthetic control of plasmid replication enables target- and self-curing of vectors and expedites genome engineering of Pseudomonas putida’, *Metabolic Engineering Communications*, 10, p. e00126. Available at: https://doi.org/10.1016/j.mec.2020.e00126.

Wong, S.M. and Mekalanos, J.J. (2000) ‘Genetic footprinting with *mariner* -based transposition in *Pseudomonas aeruginosa*’, *Proceedings of the National Academy of Sciences*, 97(18), pp. 10191–10196. Available at: https://doi.org/10.1073/pnas.97.18.10191.

Worsey, M.J. and Williams, P.A. (1975) ‘Metabolism of toluene and xylenes by Pseudomonas (putida (arvilla) mt-2: evidence for a new function of the TOL plasmid’, *Journal of Bacteriology*, 124(1), pp. 7–13. Available at: https://doi.org/10.1128/jb.124.1.7-13.1975.
